# Higher order repeat structures reflect diverging evolutionary paths in maize centromeres and knobs

**DOI:** 10.1101/2025.01.31.635908

**Authors:** Rebecca D. Piri, Mingyu Wang, M. Cinta Romay, Edward S. Buckler, R. Kelly Dawe

## Abstract

**Background:** Highly repetitive tandem repeat arrays, known as satellite DNAs, are enriched in low-recombination regions such as centromeres. Satellite arrays often contain complex internal structures called higher-order repeats (HORs), which may have functional significance. Maize is unusual in that its satellites occur in two distinct genomic contexts: centromeres, which interact with kinetochore proteins, and knobs, which undergo meiotic drive in the presence of Abnormal chromosome 10. Whether maize centromeres or knobs contain HOR patterns, and how such patterns relate to function, remains unclear.

**Results:** Here, we generate 13 repeat-sensitive genome assemblies of maize and its recent ancestor, teosinte. We develop a new graph-based pipeline, HiReNET, to classify HORs and demonstrate its utility in both Arabidopsis and maize. We find that HORs are ubiquitous in maize satellites but are typically low-frequency with small patterns, unlike the large, continuous HOR blocks characteristic of human centromeres. Approximately 38% of centromeric CentC monomers occur in HORs; however, no specific HOR class dominates any functional centromere as marked by centromeric histone H3. Arabidopsis centromeres have a similar HOR landscape. In contrast, maize knobs exhibit a more structured HOR distribution. Large knobs contain megabase-scale similarity blocks with repeated HOR patterns. These repeat units likely promote unequal crossing over, enabling rapid knob expansion, and may harbor motifs recognized by trans-acting factors involved in meiotic drive.

**Conclusions:** HORs occur in all major maize satellite arrays. Specific HORs are not associated with centromere function, but knobs contain conserved HOR patterns within similarity blocks that may facilitate meiotic drive.

## Background

Long tandem repeat arrays, called satellites, frequently occur in centromeric and subtelomeric regions of eukaryotic chromosomes. The weakened selection in these low recombination regions provide a haven for repetitive DNA, allowing it to replicate and spread with lowered risk of being purged [1]. Despite the prevalence of satellite DNAs, their origin, sequence arrangements, mechanisms of accumulation, and potential functional significance are often obscure. Recent advances in sequencing technologies have begun to reveal the internal structures of some of the longest repeat arrays in both animals and plants. In human centromeres, there is a core of alpha satellites with highly-homogenized HOR patterns that are associated with active kinetochores marked by the histone H3 variant CENP-A/CENH3 [2,3]. The HORs in core regions can differ dramatically among individuals, both in copy number and in distinct monomer patterns [4]. Outside the central core are divergent layers, representing obsolete centromere cores, that have accumulated variants and transposable elements over time. This dynamic, where highly-homogenized HOR patterns are actively evolving in close contact with the kinetochore, has been described as kinetochore selection on alpha satellite HORs [5]. A similar connection has been observed in Arabidopsis, although the HOR pattern layers and monomer variants are not as well-defined [6].

Maize also has a centromere-associated satellite called CentC [7]. However, the association between CentC and CENH3 is polymorphic – functional centromere regions can be completely decoupled from CentC arrays in some chromosomes [8–10]. Additionally, maize harbors another class of widespread satellites known as knob repeats. Knobs are large, heterochromatic satellite arrays on chromosome arms, comprising two repeats, knob180 and TR1 [11,12]. In the presence of a chromosomal variant known as abnormal chromosome 10 (Ab10), they are subject to meiotic drive. Ab10 encodes two specialized kinesin proteins (KINDR and TRKIN) and several large knobs [13,14]. During meiosis, the kinesins associate with arrays of knob repeats (KINDR with knob180 repeats, and TRKIN with TR1 repeats) and move to the spindle poles more quickly than centromeres, allowing knobs to dictate chromosome movement. As a result, when Ab10 is heterozygous, the haplotype is preferentially passed on through the female germline (up to 83%) [15,16]. Importantly, knobs on other chromosome arms can take advantage of the trans-acting kinesin proteins and show the same levels of meiotic drive when they are heterozygous [16,17]. The signature of this drive system is present in all maize genomes and their ancestors– knob repeat arrays dispersed across all chromosome arms, although only the largest arrays in mid-arm positions are known to show strong drive [16]. Knobs are defined by sequence, which is inert and densely repetitive, similar in structure to the satellite DNA associated with centromeres.

For the study of repeat arrays, maize presents an interesting test case where there are both centromeric satellites and meiotically-driven knob satellites. These repeat arrays can contribute as much as ∼500 Mb to the maize genome, though the amounts vary from line to line [13]. Recent genome assemblies have included long spans of the major maize satellites [10,18], yet there has been no comprehensive analysis of their internal makeup, or their conservation among lines. Based on the emerging data from human and Arabidopsis, we anticipated that both centromeric and knob arrays would contain HORs, and that the most homogenous arrays would occur in regions of functional relevance. Yet, given that many maize centromeres have shifted away from CentC altogether [8–10], we anticipated that the HOR patterns associated with CentC may be functionally irrelevant, and therefore might be sparse or degraded. In contrast, given that knobs are in areas of high recombination and under active selection for meiotic drive, we anticipated that the HOR patterns in knobs would be pronounced, especially in larger knobs that are more strongly driven.

To evaluate the presence of HORs, we used PacBio HiFi sequence data to analyze 13 genomes – 10 of maize and 3 of its recent, wild ancestor, teosinte. After discovering that software used to identify and describe HORs in human and Arabidopsis did not perform well in maize, we developed new methods, which use similar monomer identification and clustering methods to previous tools, such as AlphaCENTAURI and HORmon, but allow greater flexibility in pattern identification [19,20]. We found that locally-confined, small HOR patterns are widespread in maize satellites, but that they tend to be low-frequency and irregular. In centromeres, the location, density, or conservation of these patterns are not correlated to the current active centromere position. Rather, these relatively small and infrequent CentC patterns may be a signature of recurrent breakage and repair that is innate to satellite DNA.

In knobs, we found high-frequency HOR patterns that are punctuated regularly along satellite arrays, about 1 Mb apart, intermingling with the local HOR patterns. The long distance between repeating HOR patterns indicates they are likely driven by an alternative mechanism than local HOR’s – unequal crossing over. Further, we found that these patterns are shared among three of the largest knob180 arrays, indicating there is a mechanism for sharing sequences among these distinct satellite loci, possibly gene conversion or rolling-circle replication, consistent with classic models of centromere evolution [21,22]. Due to their consistency within and among knobs, we postulate that the conserved HOR patterns may represent functional units that are under selection for meiotic drive.

## Results

### Repeat-Sensitive HiFi Assemblies in Maize

We initially assessed older genome assemblies from the maize pangenome, generated with PacBio CLR (long single reads), for their utility in interpreting satellites [10]. But we discovered poor agreement between satellite content in input reads and assemblies, indicating assemblies did not accurately represent the repeats [10,23] (Additional file 1: Table S1). The older assemblies also had poor read alignment with new HiFi reads, possibly due to overpolishing with Illumina reads, relatively error-rich reads used to generate the assemblies, or misassembly (Additional file 2: Fig. S1-S3). So, we opted to use new assemblies based on HiFi reads (circular consensus reads), which are more accurate than the sequencing used in the older assemblies.

For this study, we generated assemblies for 13 inbred lines – 3 from inbred lines previously used in genetic studies (B73, B73-Ab10, and Mo17) [10,18,24], 7 diverse inbreds from public breeding programs, and 3 teosinte lines from the [25], representing ancestral variation. The final assemblies ranged from 2.186 Gb for B73 to 2.766 Gb for the teosinte inbred TIL01, with the total satellite repeat content ranging from 6.64 Mb in B73 to 300.48 Mb in TIL01 (Additional file 1: Table S2). This large range was expected, as the teosintes are known to have much larger knobs than most maize inbreds [26]. These HiFi-based assemblies have a more faithful representation of satellite DNA but there were still discrepancies between the proportion of satellite DNA in the HiFi sequence reads (CCS reads) and the assembled contigs for some inbreds (Additional file 1: Table S1, S2). One of the better assemblies was for the inbred CG108, which has similar repeat coverage in reads, contigs and the final assembly. Read alignment across the very large knobs in the CG108 assembly reveals uniform coverage with only one small area of potential misassembly (Additional file 2: Fig. S4). There were, however, gaps in all assemblies. Total gaps ranged from 46 in Mo17 (0 in satellite arrays) to 288 in Tx779 (11 in satellite arrays) (Table S3). For in-depth assessments of centromeric repeats, Mo17 was used due to its gapless arrays and availability of ChIP-seq data [18]. For knob repeats, the CG108 assembly was used due to the overall high quality and large gapless knob assemblies.

### Repeat Array Positions are Highly Conserved

Maize has four distinct classes of satellites related to centromeres: two associated with canonical centromeres, CentC (156bp), it’s primary centromere-associated repeat, and Cent4 (741 bp), a pericentromeric repeat linked to centromere 4; and two associated with knobs, knob180 (180bp) and TR1 (358bp) [7,9,11,27]. All four classes of satellite are organized into tandem repeat arrays, where multiple copies occur together in head-to-tail orientation. Previous pangenomic studies demonstrated that the major satellite array positions are well conserved within maize [10].

To identify repeat arrays within the assemblies, we first identified repeat consensus sequences using RepeatExplorer2 [28], then used the consensus sequences to query the assemblies. Within the 13 genomes used for this study, 151 distinct satellite array positions–151 on normal chromosomes, and 5 specific to the Ab10 haplotype– were identified (Figure 1a). Of these positions, 78 are private, meaning they only occur in one line, and 30 are shared among all genomes (Figure 1b). The high proportion of positional conservation is true even for small arrays that are not expected to be functional. The smallest array in a fully conserved position is only ∼1.5 kb, containing only 9 CentC monomers. Within each genome, there are a total of 71 to 113 satellite arrays, with knob180 consistently having the most arrays and Cent4 consistently having the least arrays (Figure 1c).

**Figure 1.**
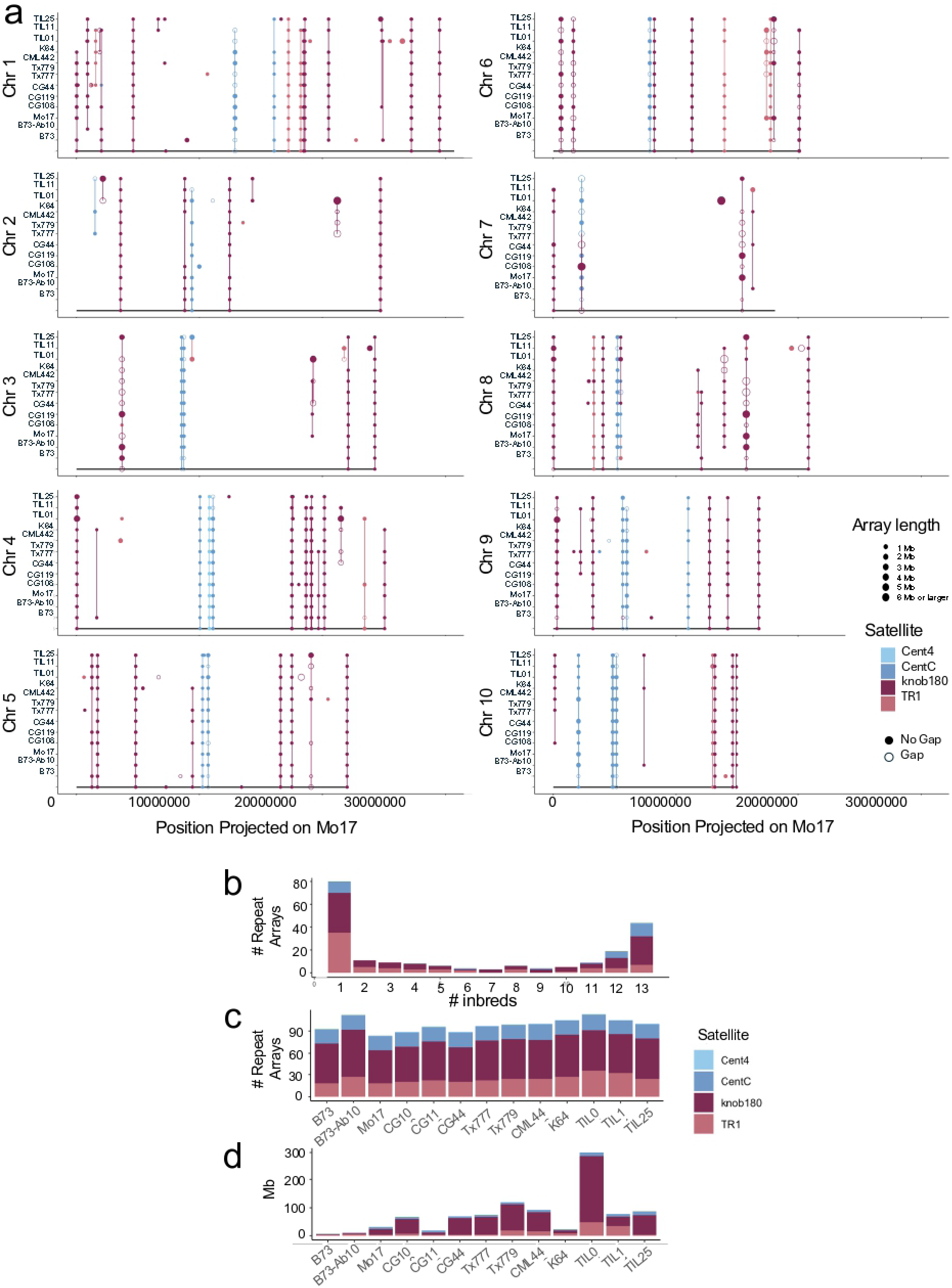
Assembly repeat content. **a**) Satellite arrays projected onto the Mo17 assembly. Open circles represent arrays that are not fully assembled (contain an N gap). Filled circles represent fully assembled arrays (do not contain an N gap). Lines connecting arrays represent homology based on synteny with conserved genes. Lines connect syntenic arrays. Single circles represent private arrays, though not all are visible because circles representing large arrays sometimes overlap smaller, closely linked arrays. The unique knobs on the Ab10 meiotic drive haplotype are not displayed on this graph. **b**) Frequency spectrum of arrays. **b**) Number of different repeat arrays by satellite. **c**) Repeat array lengths.

Although there is some variation in the number of arrays per genome, variation in array number is not sufficient to account for the overall variation in satellite content. Rather, a small number of very large arrays drive overall satellite content variation (Figure 1b-d). The variation in length at one site can be substantial. At one CentC array position on chr10, the largest centromeric array assembled without gaps is 5.7 Mb in Mo17 and the smallest is 7.7 kb in B73 – a length difference of ∼5 Mb, and a relative difference of >700x. In the largest knobs, size can vary >46 Mb at a single locus– for example, one of the largest gapless arrays on chromosome 7 is 51 Mb in CG119 but its homolog in CG44 is only ∼9% of its size at 4.8 Mb.

### Local HOR Classification

Methods for HOR identification can be categorized into two groups: first, algorithms that use high-copy, periodic k-mers to identify novel repeats and HORs de novo (i.e. TRASH, SRF toolkit); and second, methods that utilize previously-characterized satellite sequences to identify patterns of monomer subtypes (e.g. AlphaCENTAURI, HORmon, HiCAT) [19,20,29–31][19,20,23,29–31]. Current software lacks power to identify HOR patterns that are low-frequency and irregular, like those in maize. SRF toolkit is the only current algorithm that was benchmarked with maize, and it only identified two major satellite repeats in B73 – a CentC variant and a 4-mer knob180 unit, neither of which formed repeat arrays [23].

To improve HOR identification in maize, we adapted methods from algorithms that identify monomer subtype patterns. But, to better identify small, locally-confined patterns, only single 10 kb regions (bins) were analyzed at a time (Figure 2). Additionally, expectations of periodicity of the pattern, meaning that the same sequence motif recurs in a regular pattern with even spacing among all occurrences, were removed. Within a single bin, monomers were identified with HMMER [32], a method that detects truncated and degenerate monomer forms. The monomers within a bin were then compared all to all with BLAT [30,33]. Similarity between monomer sequences was calculated as a Jaccard Index (# identical bp/(total length of both sequences - # identical bp)), which represents both sequence and length similarity. Pairwise similarity scores were used to generate networks, with nodes representing distinct monomer sequences and edges present between two nodes representing similarity. Networks were regenerated for all Jaccard similarity scores between .90 to .99 in .01 increments, where the nodes remained the same, but similarity cutoffs to determine edges became progressively more stringent, similar to the processes of alphaCENTAURI, HORmon, and HiCAT [19,20,31]. We found that Jaccard scores lower than .90 led to collapse of the monomers into a single network even when HOR patterns were present (Additional file 2: Fig. S5).

**Figure 2.**
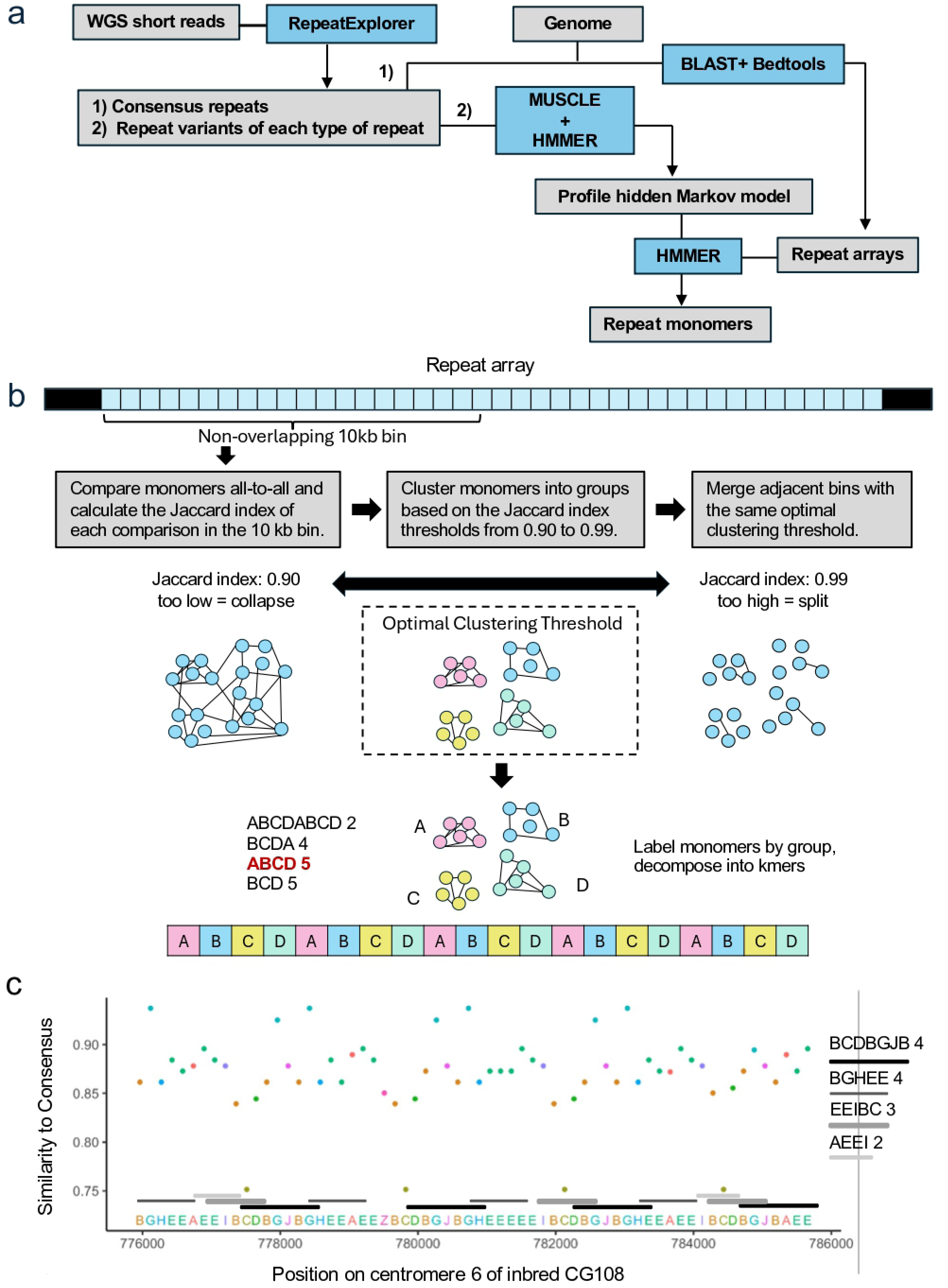
Local HOR identification. **a)** Workflow for identifying repeat monomers. Whole-genome sequencing short reads are analyzed with RepeatExplorer-TAREAN, which produces consensus repeat monomers together with their variant subtypes. The variant subtypes that capture sequence diversity that improves repeat identification by HMMER. **b**) Monomers from 10 kb bins are then compared to all other monomers in the same bin, and a Jaccard similarity index calculated for each pairwise comparison (the Jaccard index summarizes both sequence polymorphism and length differences). The output is a matrix of Jaccard similarity scores for each 10 kb bin. Then, similarity networks are generated for thresholds from .90 to .99, in increments of .01. Each node represents a monomer sequence. If two monomer sequences are at least as similar as the threshold, monomers are connected with an edge. Monomers from each distinct cluster in the network are labeled with a letter. Private clusters, or clusters that only contain a single monomer, are labeled with “Z”. Adjacent bins with the same clustering threshold are merged. Monomers, with labels of their cluster identity, are then put back in their original genomic order, resulting in a character string. The character string is decomposed into k-mers, starting with k=3 until all k-mers have a frequency of 1. K-mers are then filtered to remove larger k-mers that occur less frequently than its subset k-mers, smaller k-mers that occur as frequently as larger k-mers containing it, and k-mers that contain the private monomer “Z”. In the example given, ABCD is the largest most abundant HOR. **c**) Sample HOR region, where monomers are labeled and colored by cluster identity. HOR k-mers and their frequencies are listed on the side, and each repeated pattern in the arrays is marked with a line corresponding to its k-mer.

The repeat structure for the bin was then predicted for all thresholds with an LDA model, using summary information describing the network structure. Important network summary statistics included the following: proportion of monomers in the largest cluster (# monomers in largest cluster / # of total monomers), proportion of monomers in the second largest cluster (# monomers in second largest cluster / # of total monomers), number of unconnected clusters relative to number of monomers (number of monomers that do not share similarity with any other monomers / # of total monomers), average pairwise Jaccard Index, and proportion of monomers collapsed into the most prevalent subtype (or most common distinct sequence) (# monomers identical to most common sequence / # of total monomers). Utilizing graph structure to identify optimal clustering thresholds and predict HOR structure is novel to this study.

The LDA model classified the repeat structures in each bin at every threshold in one of three categories: HOR, where most monomers belong to two or more similarly-sized clusters in the network; Order, where most monomers are connected together in a single cluster; and Disorder, where most monomers do not belong to clusters. For each bin, the similarity threshold classified with the highest posterior probability was used for further analysis, deemed the “optimal clustering threshold”, and adjacent bins with consistent classifications and thresholds were merged (this merging step results in some bins being longer than 10 kb). For bins with predicted HOR structures, monomers were labeled by their cluster identity. For example, all the monomers in the largest cluster were labeled as “A”, monomers from the second largest cluster were labeled as “B”, and so on using both upper and lower case characters (A-Y, a-y) and numbers 0-9, if needed. Monomers that were unconnected were labeled at “Z”, representing a private cluster. Monomer patterns could then be represented as character strings, with each monomer represented by its corresponding cluster character. The pattern string was then decomposed into k-mers of various lengths to identify repeating patterns of >=3 monomer subtypes (Figure 2c).

The results showed differences in total HOR content among repeat types and inbreds, with the highest HOR percentages being found in Cent4 arrays (as high as 77%) and the lowest in knob180 arrays (as low as 6%) (Figure 2a). CentC satellites contain slightly more HOR content (38%) than knob180 arrays (28%) but this difference is not statistically significant when considering the variation among inbreds (p=0.065, two-tailed t-test). While HORs are common in maize repeat arrays, there are thousands of different patterns, and each HOR is usually composed of only a few monomers and occurs at a low frequency (Figure 3b, 3c). The vast majority of HORs are confined to a single bin on an array. When an HOR occurs in more than one bin on the same centromere it is usually found in only 2-3 bins (Figure 4 and Table S4; the methodology for identifying shared HORs is described below).

**Figure 3.**
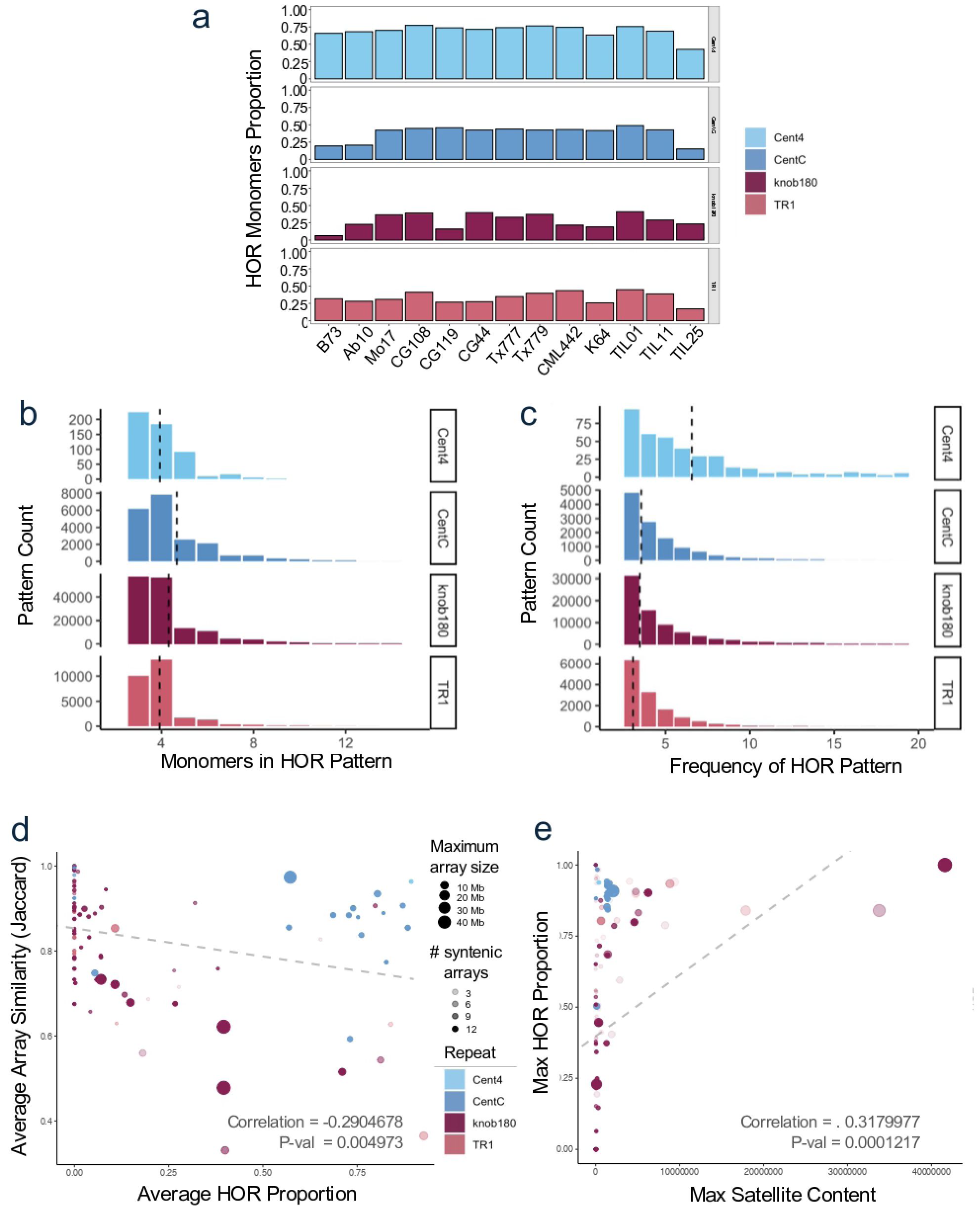
HOR Pattern Frequency and Length. **a**) Proportion of HOR monomers identified for each line. **b)** HOR pattern length distribution. **c)** HOR pattern frequency distribution. Dashed line marks mean **d)** The maximum HOR proportion is negatively correlated with average Jaccard similarity of arrays at syntenic positions **e)**. The maximum satellite content is positively correlated the maximum HOR proportion of the arrays. Only gapless arrays are plotted in (d) and (e); linear fits to these data were significant (p < 0.01).

**Figure 4.**
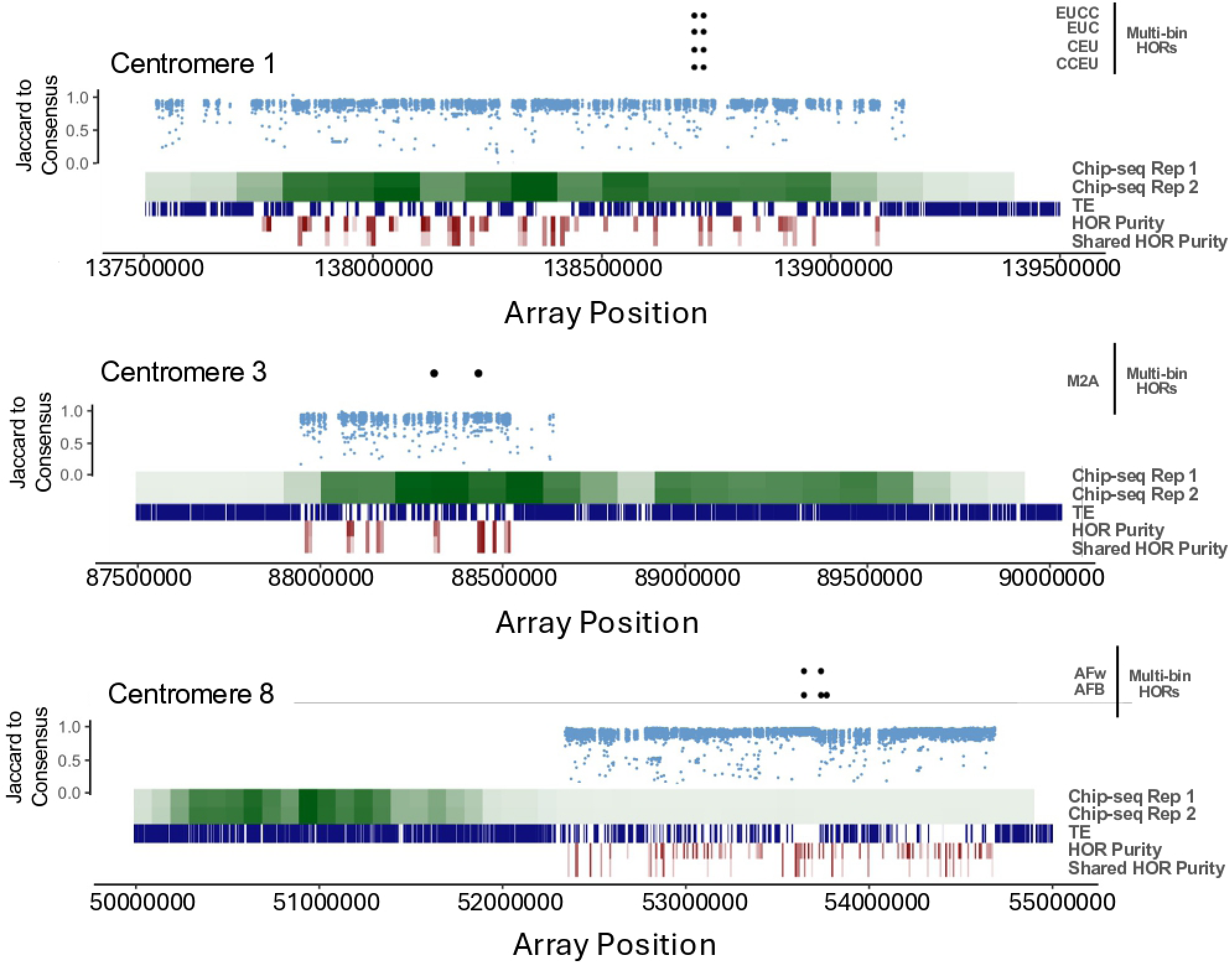
CENH3 Chip-seq on Mo17. For each centromere, multi-bin HOR patterns that occur in at least two distinct HOR regions are indicated in the upper panels, where dots indicate presence in the array. The Jaccard similarity of each monomer to the consensus is shown as a dot plot. The relative CENH3 ChIP-seq density in 100kb bins (of two independent replicates) is shown in shades of green. TEs are represented by blue boxes. HOR purity and shared HOR purity within bins are shown in red, where darker shades represent higher purity values.

To enhance accessibility and usability, we streamlined our pipeline into a downloadable tool called *Higher-order Repeat Network Exploration Tool* (HiReNET) and applied it to the Arabidopsis ecotype Ey15-2 [34]. We chose this ecotype because centromere 1 was featured in the published article as having a discrete region with a particularly high “HOR score” (from ∼16.3 MB to 17.6 Mb, Additional file 2: Fig. S6). The HOR score is a repetitiveness measure that describes how often a monomer in an HOR is likely to be found in any other HOR [29,34]. The results of running HiRiNET on Ey15-2 showed that most HOR patterns are 3-5 monomers in size, in agreement with earlier conclusions [34]. Arabidopsis ecotype Ey15-2 has fewer HORs than maize Mo17 (29% of all monomers are in HORs) but has more multi-bin HORs (Table S4). In the ∼1.3 Mb central region of centromere 1 denoted as having a higher HOR score, there was no obvious enrichment of HORs, though the optimal clustering threshold for HORs in this region was notably higher (Additional file 2: Fig. S6). Some of the lower-threshold multi-bin HORs occur on both sides of this central zone, supporting the interpretation that active repeat turnover in the center pushes older repeats to the edges [3,6].

### Syntenic Arrays Share Sequence Similarity

Syntenic arrays, which are conserved in positions relative to core genes, are also generally conserved in sequence. We compared entire satellite arrays in a manner similar to our monomer comparisons, by calculating the total sequence identity between syntenic arrays, excluding any non-satellite DNA or TEs, then computed a Jaccard index ((# identical bp/(total bp of both arrays - # identical bp)). The syntenic arrays have an average Jaccard similarity of .82 (Table S5). This similarity score captures both sequence and length divergence and varies slightly among satellites– ranging from average similarity of .96 for Cent4 positions to .74 for TR1 positions. Comparing two non-homologous arrays will generally give much lower values: for instance, aligning the knob on chromosome 3 from the inbred K64 (33398 bp of knob180 repeat) to the knob on chromosome 9 from K64 (183257 bp of knob 180 repeat) give a match of 26487 bp and a Jaccard index of 0.14.

Within a group of syntenic arrays, smaller arrays are generally more similar to each other than larger arrays (Figure 3d). Smaller arrays also have less relative HOR content (Figure 3e). In fact, the smallest arrays, where the largest homolog is <=10kb, are completely devoid of HORs. Larger repeat arrays have higher proportions of HOR content and lower average similarity among syntenic arrays, suggesting that greater HOR content is related to greater divergence (Figure 3d,e). This trend is more pronounced for knob180 arrays, and likely reflects a tendency for longer arrays to undergo rapid expansion and contraction events.

### Shared HOR Classification

Patterns were compared among 10 kb bins to capture HORs beyond local regions, including non-adjacent bins on the same array and those in syntenic arrays in other inbreds (Figure 5). To do this, all monomers from HORs were converted to consensus sequences (i.e. pattern ABCABC was converted to consensus monomers A, B, and C). Then, consensus monomers were compared all-to-all. A network was created, where each node represents a single consensus monomer, and two nodes are connected by an edge if their sequences are at least as similar as their shared optimal clustering threshold. Meaning, if patterns ABC and DEF with optimal clustering threshold of .95 are being compared, A, B, and C are expected to cluster separately at the .95 cut off. However, if these patterns share recent evolutionary history, A may cluster with D, B with E, and C with F. This step is necessary since local HORs were initially labeled based only on patterns within their local 10 kb bin. By reclustering, we could relabel HORs based on larger-scale comparisons and generate a key to translate among regions.

**Figure 5.**
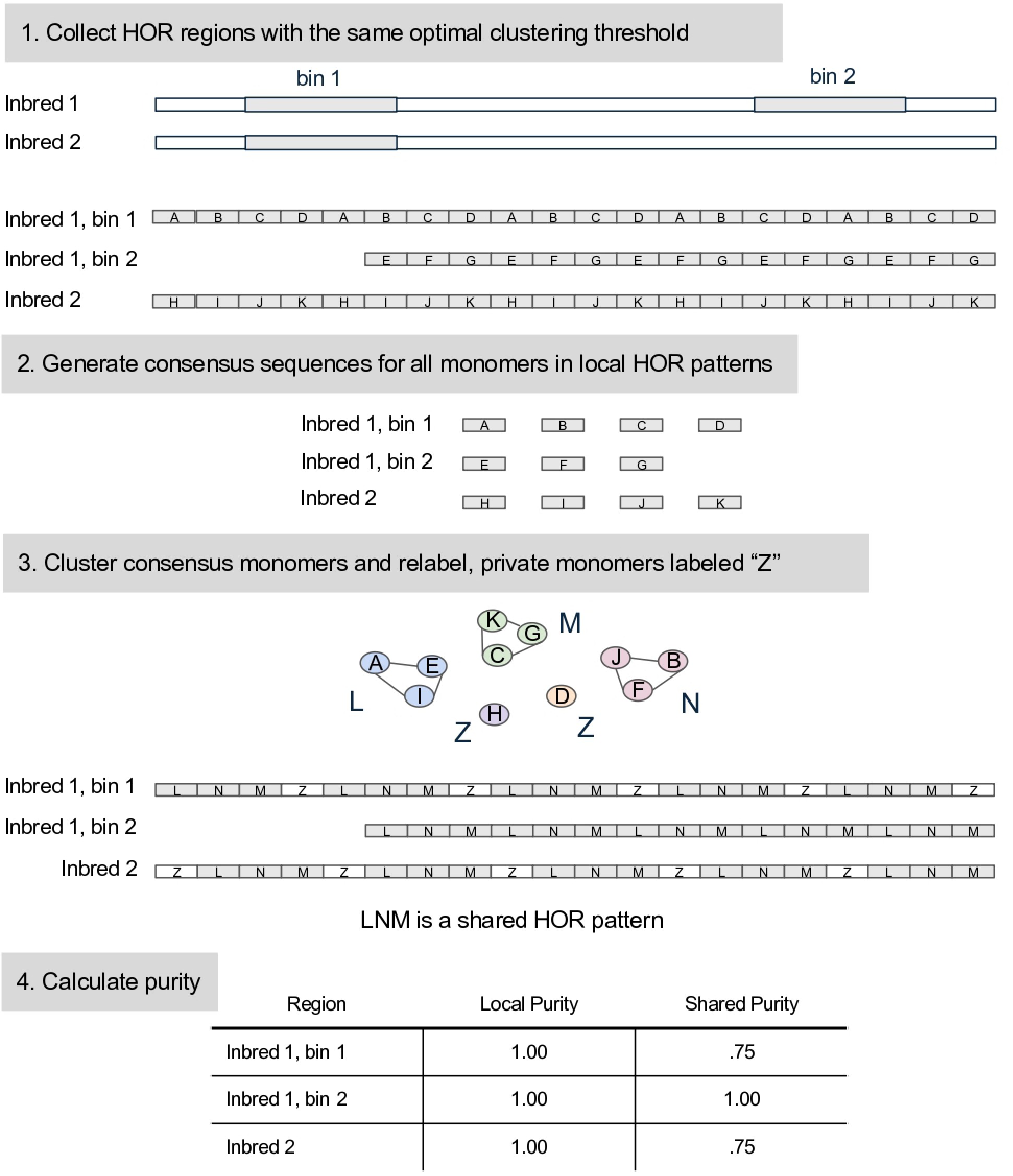
Shared HOR Identification**. 1**. HORs with the same optimal clustering thresholds are extracted. This can include multiple distinct regions from the same array. **2**. Consensus sequences for each monomer subtype in an original HOR pattern. **3**. Consensus sequences are then compared all-to-all with a cutoff equal to the original shared optimal clustering threshold, and a network is generated. Monomers are labeled by their cluster identity and returned to their original orders. Kmers are then generated. Only kmers that exist in one or more regions are shared HORs. **4**. Array purity is recalculated as (N monomers in identifiable arrays) /(total monomers).

To facilitate display of these data we used a metric we refer to as HOR purity, which is the number of monomers that occur within HORs divided by the total number of monomers in a bin. An extension of this metric is shared HOR purity, which is a recalculation of HOR purity after including a conserved syntenic region (Figure 4, S7, S8). Of all HORs in Mo17, our model assembly for centromeres, only 14% of the HOR patterns within an array are also observed in the syntenic array of at least one of the other 12 genomes (Table S6). We found that 13% of the HORs in CentC arrays, 88% of the HORs in Cent4 arrays, 8% of the HORs in knob180 arrays, and 62% of the HORs in TR1 arrays are observed in syntenic arrays of at least one inbred. Similar results were observed for CG108, our model assembly for knobs (Table S6). The data suggest that CentC and knob180, the more prevalent satellites that are related to active centromeric and neocentromeric function, are undergoing more rapid change, which leads to less overall relatedness among syntenic arrays.

### Centromeric HORs are not visibly enriched in CENH3 domains

Of the 13 maize lines used here for genome analysis, CENH3 ChIP-seq data are available for only one, Mo17 [18]. CENH3 centers around the major CentC array on a subset of the chromosomes (1, 4, 7, and 9), is partially associated with CentC on other chromosomes (2, 3, 6 and 10), and is completely decoupled from CentC on others (5 and 8) (Figure 4, S7, S8) [18]. The fact that CENH3 domains can be decoupled from nearby CentC arrays is well documented [8–10] and our data confirm the conclusions of the prior Mo17 study [18]. A visual inspection of these centromeres reveals no clear correlation between HORs and CENH3 localization. This is particularly evident in the comparison of Centromere 1 and 8 (Figure 4). On Centromere 1, where CENH3 is well centered over the CentC array, the HOR percentage (monomers in HORs/total monomers) HORs is 39%, whereas on Centromere 8, where the CENH3 domain is entirely displaced from the CentC array, the HOR percentage is 41% (Table S4; the HOR percentages for all Mo17 centromeres are normally distributed and both the 39% and 41% values fall within one standard deviation of the mean).

### Knobs Contain Conserved Patterns

Knob sizes are known to vary greatly among maize lines [35], and have the capacity to dramatically increase or decrease in size as a result of genetic exchange events [36]. We wondered if there might be any unique features within knobs that might enable rapid changes in size. Visualization of plots showing the Jaccard similarity of knob180 monomers to the consensus sequence revealed clear megabase-scale periodicities. These large-scale patterns, roughly .48-1.5 Mb, are defined by regions of high similarity to the knob180 consensus followed by regions with low similarity to the consensus, visible as vertical stripes of low-similarity monomers (Figure 6). We refer to the units of the large-scale patterns as similarity blocks. Within a single similarity block, the regions of low homology to the knob consensus have degraded knob repeats, while the regions of high consensus have more intact arrays and HORs. When 1 Mb units of these patterns are aligned to each other in a traditional all-by-all dot plot, there is strong homology along the diagonal, suggesting that they are related by descent (Figure 7).

**Figure 6.**
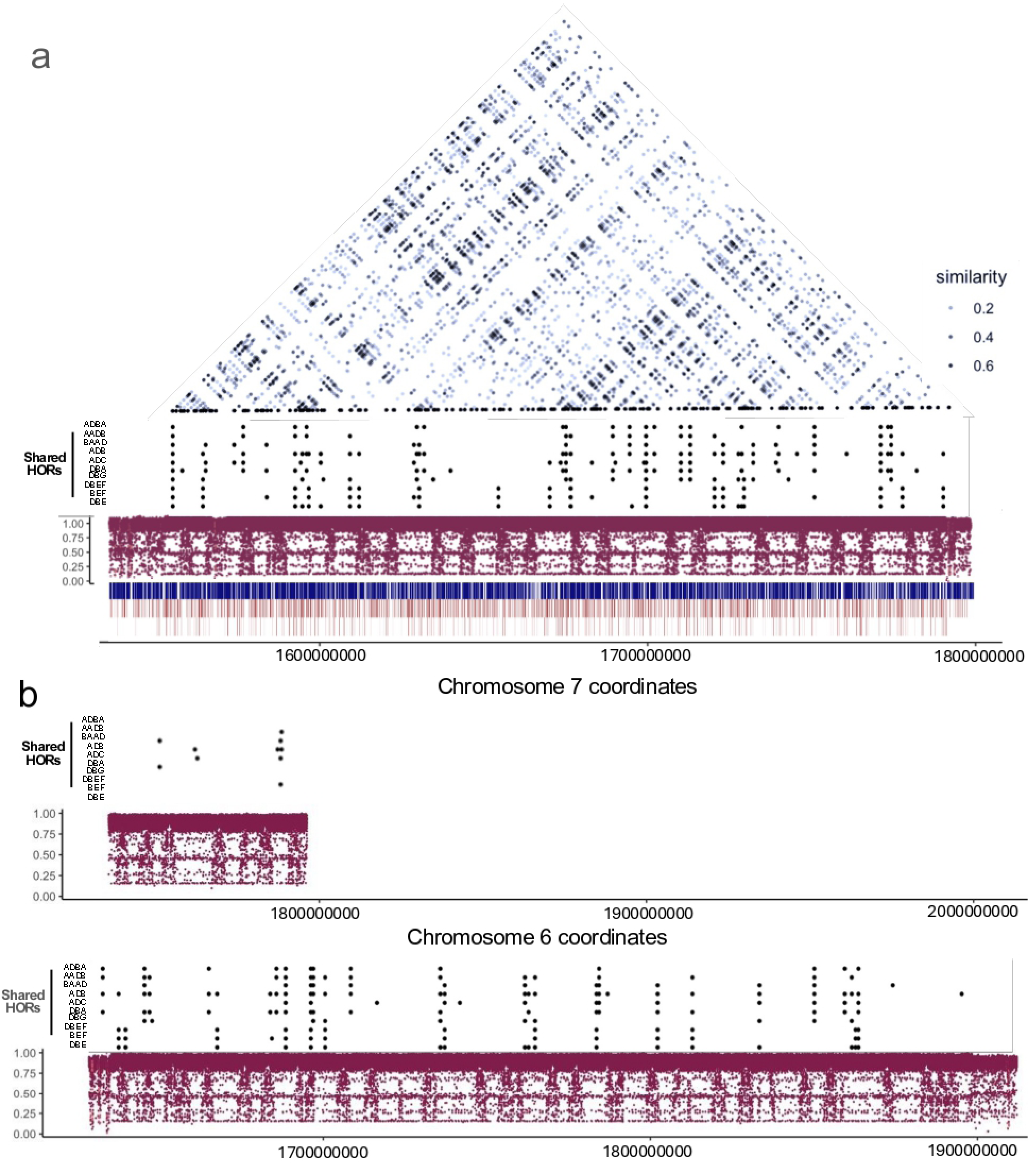
Shared HORs among knob180 arrays. **a**) The large knob on chromosome 7L in CG108. The upper triangle shows HOR pattern similarity of all HOR regions of the array. Darker shades indicate a greater proportion of shared patterns, up to 1. Below the triangle are high frequency HORs that are shared among all three knobs (including those in b), where dots indicate presence in array, and the x axis is genomic position. Below the shared HORs, in maroon, is a dot plot of monomers displayed according to similarity to the consensus as a Jaccard index on the y axis. Each maroon dot represents a monomer. The heat maps, from upper to lower, show TEs (blue), HOR purity (red), and shared HOR purity (red) (same labeling as Figure 5). **b**) The knobs on chromosome 6L and 8L in CG108. Each panel shows high frequency shared HORs found in all three knobs, and dot plots of monomers displayed according to similarity to the consensus as a Jaccard index.

**Figure 7.**
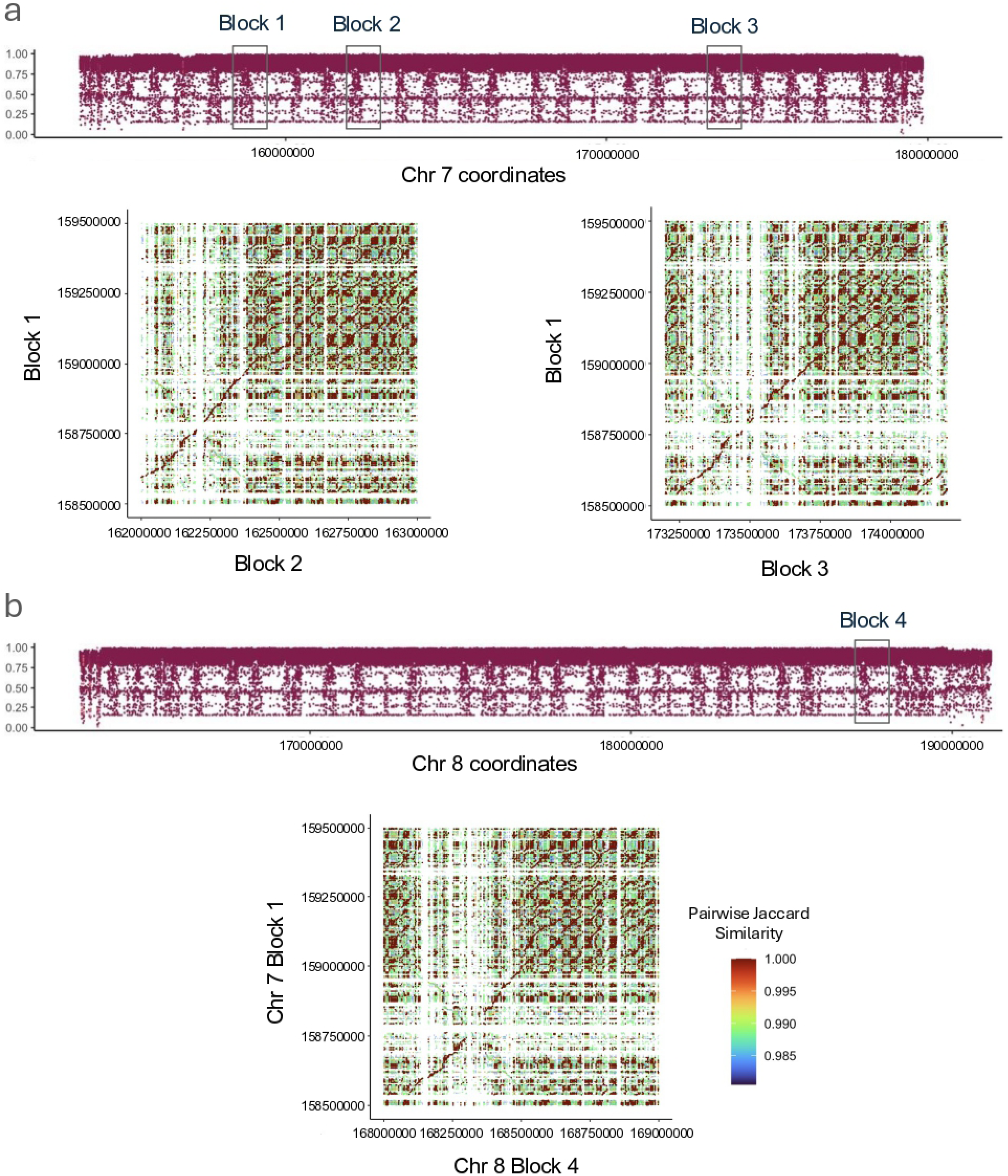
Similarity Blocks in knobs. **a**) Similarity blocks in the large knob on chromosome 7L in CG108. The maroon similarity to consensus plot (same plot as Figure 6a) shows the locations of three similarity blocks identified by their characteristic features of having a region of low similarity to the consensus followed by a region of high similarity to the consensus on a megabase scale. Below, there are two traditional dot plots comparing blocks 1 and 2 and blocks 1 and 3. Note the diagonal lines of high similarity. **b**) A similarity block in the large knob on chromosome 8L in CG108 (labeled block 4). Below is a dot plot comparing chromosome 7 block 1 to chromosome 8 block 4. Note again the diagonal line of high similarity.

The similarity blocks are also typified by many multi-bin HOR patterns. In the fully assembled knob on the long arm of chromosome 7 in CG108, there are 417 distinct HOR patterns that occur in more than one similarity block, six of which occur in at least 30 blocks across the array (Figure 6a). These shared patterns exhibit a skipping pattern, where the conserved patterns are spread across the array in regular intervals. The same shared HOR patterns are found on the other two large knobs in CG108, which are on chromosomes 6 (6Mb) and 8 (28Mb) (Figure 6b). There are 677 HOR patterns that are shared among at least two of the knobs. The 10 highest-frequency, shared HOR patterns are composed of 7 high-frequency monomers, which are named here as knob180 A-F (Table S7, S8).

## Discussion

In this study, we developed a novel HOR identification pipeline (HiReNET) that revealed that a large proportion of satellite content in maize, both in centromeres and knobs, are composed of higher-order repeat patterns. Unlike HOR patterns previously described in humans [3], maize HORs are primarily locally-confined, meaning most occurrences of a single HOR pattern are contained within a region of ∼10 kb. Applying HiReNET to Arabidopsis revealed a similar centromere architecture where most HOR patterns are local and rarely reiterated across multiple bins. In both maize and Arabidopsis, HOR patterns are small and low-frequency– with average pattern sizes of ∼4 monomers with ∼3 occurrences.

While pervasive in the satellite space, the presence of HOR patterns does not seem to be related to active centromere activity in maize. In human centromeres, there are older layers of HOR patterns that are present on either side of the centromere core. Drawing a line to connect regions with shared HOR patterns, human centromeres look like a layered onion [3]. The limited Arabidopsis data we generated suggests that some Arabidopsis centromeres may also have the outlines of an onion-like arrangement (Additional file 2: Fig. S6; [34]). Such layers are absent in maize centromeres. While shared patterns among regions of the same array exist in some maize centromeres, they do not exhibit any obvious patterns (Figure 4, S8). Meaning, two bins with shared patterns are near each other, rather than existing on opposite sides of the array. These results suggest that maize centromeric satellites undergo different selection than human centromeric satellites, which have evidence of kinetochore selection on pure HOR patterns [5].

The presiding evolutionary model for the origin and maintenance of HOR patterns is breakage induced replication (BIR), which proposes that repeats take advantage of the inherent instability and breakage at centromeres, co-opting repair mechanisms to expand existing patterns, generate new repeat variants, and exchange repeats among different arrays or chromosomes [37–39]. Under this model, HOR pattern expansions are frequent but relatively small, as strand repair is expected to be primarily over local distance (i.e. on the scale of a single HOR pattern length, upwards of a few dozen monomers). Experimentally, this model is supported by HOR copy number changes over 20 somatic cell divisions in a sensitized human cell line, but the frequency of the events within an organismal germ line is unknown [40]. This model unifies evidence of replication fork stalling and collapse at centromeres (reviewed in [37]) and high levels of centromeric rearrangements and instability [41–43] with observed repeat structures, as well as proposes a feasible colonization mechanism across different arrays via BIR. The local HOR patterns in maize centromeres, which are only present in longer arrays, seem to suit expectations that late-replicating, highly repetitive regions of the genome are enriched for DSBs, which can be repaired out-of-register and result in short-range tandem duplications [41,44]. However, there is no evidence that the kinetochore applies selection for enriched BIR in centromeric cores.

On the other hand, knobs do have signatures of selection. We have described large-scale patterns of similarity in some of the largest knobs that contain conserved HOR patterns (Figure 6, 7). The shared HOR patterns among knob regions are dispersed at semi- regular intervals, punctuated with local patterns in between – like skips across the array.

Mechanistically, the scale of these long-range skips seems unlikely to be driven by BIR due to the large distance between the block patterns. Rather, because knobs are located on chromosome arms where recombination is high [45], unequal crossover may be a more apt explanation [46]. The unequal crossover model is based on classic literature showing that two tandemly duplicated genes can either contract to one copy or expand to three copies as a direct result of crossing over [47,48]. Although there are no known genes in knobs, there is evidence that repetitive knob DNA pairs during meiosis and undergoes crossing over. Stack and colleagues demonstrated that foci of staining for MLH1, a key protein required for meiotic crossover [49], regularly occur within paired knobs at the pachytene substage of meiosis [50]. During pairing in repeat-dense regions, alignment can easily slip out-of-register and result in unequal crossing over. In the largest knobs, misalignment of similarity blocks could occur on a megabase scale, resulting in major shifts in total knob size [36]. Therefore, within knobs unequal crossing over may drive large expansion events, while BIR drives an abundance of smaller-scale, local HOR patterns. Drive may positively select on both expansion types, as both mechanisms can increase the array sizes (although on different scales), as well as select for sequences specific to the high-frequency monomers.

In lines carrying abnormal chromosome 10, KINDR associates with knob180 knobs to initiate neocentromeric activity during cell division [13]. Larger knobs are known to drive better than smaller knobs, possibly indicating better binding affinity to the protein [51]. While the binding specificity of KINDR to knob repeats is well documented, how that interaction is achieved is unknown. An unknown linker protein is hypothesized to interact directly with the DNA sequence and recruit KINDR [16]. The distribution of the conserved knob repeats and HORs containing those repeats leads us to believe that they may function as the binding sites for the postulated linker protein. Ongoing selection on these sequences for meiotic drive would also explain how they maintain their identity and structure over such long distances in multiple loci.

### Conclusions

We have demonstrated that the maize satellite landscape is replete with low-frequency, low-periodicity HORs that are not detectable using approaches developed for other model organisms (e.g [52–55]). These HOR patterns are consistent with random breakage and repair by BIR. Unlike in humans [3], there is no apparent enrichment of HORs in regions occupied by kinetochores, suggesting little if any functional relationship between HORs and centromere function. In contrast, knobs have repeat structures suggestive of functions related to meiotic drive. There are large, megabase-scale similarity blocks that may facilitate rapid expansion and contraction of knob size by unequal crossing over. Within the similarity blocks are conserved HOR patterns that may serve as binding sites for meiotic drive proteins.

## Methods

### Data Sources

Raw reads from the maize NAM pangenome resource [10], including Illumina and PacBio CLR data, were retrieved from PRJEB31061 and PRJEB32225. Maize pangenome assemblies were downloaded from MaizeGDB.org. Raw reads, including PacBio CLR, Illumina, and ONT, and the previous assembly of B73-Ab10 were collected from PRJEB35367. PacBio HiFI data for B73-Ab10 [56] was obtained from PRJNA1254310. Raw PacBio HiFi reads from Mo17 were collected from PRJNA751841 [18]. Raw PacBio HiFi reads for B73 were collected from SRR11606869 [57]. Raw PacBio HiFi reads for TIL01, TIL11, and TIL25 were collected from Bioproject PRJEB50280 [25].

For all other assemblies, DNA extraction and genome sequencing was performed at the Arizona Genomics Institute (The University of Arizona). Genomic DNA was extracted with a modified CTAB method [58]. High molecular-weight DNA was quality checked with Qubit HS (Invitrogen) and Femto Pulse Systems (Agilent) and 10 µg DNA were sheared to appropriate size range (15-20 kb) using Megaruptor 3 (Diagenode). PacBio HiFi sequencing libraries were constructed using SMRTbell Express Template Prep Kit 2.0. The library was size-selected on a Pippin HT (Sage Science) using the S1 marker with a cutoff at 15 kb. The sequencing libraries were sequenced on a PacBio Sequel IIe instrument with PacBio Sequel II Sequencing kit 2.0.

Raw PacBio HiFi reads for CG44, CG119, CG108, Tx777, and Tx779 are available under NCBI Bioproject PRJEB59044, and were sequenced as part of the Genomes to Fields Project. Raw PacBio HiFi reads for K64 and CML442 are available under NCBI Bioproject PRJEB66502 and were sequenced as part of an effort to sample diverse maize haplotypes containing sources of unique alleles available at the USDA-ARS maize Germplasm Resources Information Network (GRIN).

### Repeat Consensus Sequences

To capture repeat content and variation, de novo repeat identification was performed with RepeatExplorer2 (v3.6.4) (-p -put ILLUMINA -c 20 --max_memory 500000000 -tax VIRIDIPLANTAE3.0) using PE150 Illumina data for NC350 from the maize pangenome [10,28]. RepeatExplorer uses TAREAN under-the-hood to reconstruct repetitive DNA through de Bruijn graphs for high-copy k-mers. The output provides both a single primary consensus, based on the best-supported graph path through the most frequent k-mers, and a list of common variants in the data, based on the other common k-mers. The primary consensus sequence and consensus variants for the satellites of interest (CentC, Cent4, knob180, and TR1) were identified in the RepeatExplorer output by comparing sequences to previously described consensus monomers using BLAST+ [9,59].

#### Identifying repeats in reads and assemblies

Using only a single primary consensus sequence can be an issue in tandem repeat analysis, as blast prioritizes maximum similarity to a single sequence at a time, resulting in many overlaps in the output data and partial copies [60]. In decomposing a repetitive region into individual monomers, finding distinct monomers is key for downstream structural analysis [30]. As an alternative to using only a single consensus and blast for finding full-length monomers, nhmmer was used. Consensus variants identified by RepeatExplorer were aligned with a multiple sequence alignment in MUSCLE (v3.8.1551) and used to generate phmm’s (profile hidden markov models) with the makehmmerdb command in HMMER (v3.3.2) using default parameters [32]. Rather than reporting all hits, including overlapping and partial hits like BLAST, HMMER utilizes a collection of sequences within one phmm to find the best match for each region. Only the best, most complete similarity hit for each monomer is reported when repeat monomers are identified using the nhmmer command.

### Repeat Content

To calculate estimated total repeat content, satellite consensus sequences were compared to raw reads of multiple classes and genome assemblies using Blast+. For TE enrichment analysis in raw reads, the analysis was repeated using Shojun Ou’s TE library [61]. Repeat hits were converted to a bed file and then merged using bedtools (v2.3) [62,63]. Total length of reads and assemblies was counted with Bioawk (v1.0) [64]. Total genomic proportion of repeats was then calculated by dividing total repeat length, summed from merged blast hits, by total read (or genome) length. Alternatively, estimated genomic content was determined by dividing the total repeat length by the estimated read depth. Estimated repeat depth for HiFi reads was reported by hifiasm [65]. For other read types, estimated read depth was gathered from their source papers [10,24].

### Read Depth Over Repeats in Old Assemblies

Raw PacBio CLR reads from the NAM pangenome project [10] were aligned to the B73-Ab10 assembly and two maize pangenome assemblies (B73, NC350) to check read depth over centromeric regions [10,24]. Reads were aligned to both full assemblies (with unscaffolded contigs included) and only the pseudomolecules (representing only assembled chromosomes). For the B73-Ab10, ONT and HiFi reads were also used. In each case, reads were aligned with minimap2 (v.2.24) with appropriate default settings for the data type [66,67]. Read alignments were filtered using the 2308 filter with samtools (v1.6) [68] to remove multi-mapping. Alignment bam file was then converted to a bed file using bedtools (v2.3) bamtobed with default parameters, with a fourth column of read length added. Average read depth over 10 kb was calculated with bedtools map (v2.3) (-c 4 -o sum) for each alignment [62,63] (Quinlan and Hall 2010). For each bin, total read length was summed and divided by 10 kb. Bin content of satellites was also identified, using bedtools intersect to combine satellite blast hits with 10 kb bins. Satellite hits for each bin were summed and divided by 10 kb to identify repeat proportion.

### HiFi Assembly

HiFi contigs were generated using hifiiasm (v 0.19.6) with homozygous settings with end-joining disabled (--write-ec --write-paf -u0 -l0) [65]. Contigs were checked for repeat content using BLAST+ (v2.10.1) (blastn -outfmt 6 -num_threads 10 -max_target_seqs 5000000) [59]. The resulting contigs were variable in quality– for many genomes, the contig-level assembly was enriched for short, low support contigs that were dense with satellite content (Additional file 2: Fig. S3b). The presence of these contigs was unrelated to read depth, length, or quality. These low-confidence contigs inflated the total relative repeat content, with contig repeat content outpacing genome estimates from raw reads. To enrich for accurate array assemblies, low-confidence contigs were removed, leaving only contigs with greater than half the expected read support (i.e. for an inbred with HiFi read depth of 20, only contigs with >10 read support were used). The high-confidence contigs are longer, have repeat content consistent with raw reads, and are largely anchored, meaning they have at least one edge that is not satellite DNA and can be confidently scaffolded into a final assembly. High confidence contigs were scaffolded using the Mo17 assembly using RagTag (v2.0.1) [18,69]. To identify anchored repeat arrays for further study, we used bedtools intersect to identify contigs that had repeats within 100 bp of one end a contig. The assembled CG108 genome was validated for heterozygosity using NucFreq, which estimates primary and secondary allele coverage from sequencing reads [70].

In the initial assembly for CG119 scaffolded with Mo17, chromosome 3 had two large CentC arrays, and chromosome 7 had none. The CG119 assembly scaffolded with B73 v5 did not have this same anomaly. To manually correct this issue, the misplaced contig was manually added to a bed file in the correct spot for chromosome 7 and removed from chromosome 3 in another bed file. 100N gaps were placed between contigs.

### Annotation of Core Genes and Use in Identifying Syntenic Array Positions

Scaffolded assemblies were annotated with Liftoff (v1.6.3), using gene annotations from Mo17 as the reference [71]. Annotated genes were subset down to a highly conserved set, defined as genes that are single-copy in all new assemblies, mapping to the same chromosome, and were core genes in the maize pangenome [72]. To identify core genes, gene coordinates were collected from the annotation GFF file and converted to a bed file. Then, gene annotations for B73 v5 were similarity converted, subset to the list of known core genes, and extracted as a fasta file using bedtools getfasta (v.2.29.2). The B73-derived core genes were then aligned to Mo17 with minimap2 (v2.22), filtered to their best hit using samtools view (-F 2308) (v.0.1.2), and converted to a bed file using bedtools bamtobed [66–68]. B73 core gene hits were then intersected with Mo17 annotations to find equivalent genes with bedtools intersect.

Repeat monomers previously identified via BLAST+ were filtered to a minimum size of 30bp and merged within 10kb with bedtools to define arrays. Array coordinates were compared to the core genes using bedtools to identify the nearest upstream gene (-iu -D a -a) and downstream gene (-id -D a -a), which were used as the array coordinates for comparison. Array positions were clustered among lines using the graph_from_edgelist function in igraph (v1.2.6), with each node representing an array, and edges among arrays indicating that the two arrays share either or both core genes on either side [73], [74]. Repeat arrays in the same position relative to conserved genes were referred to as syntenic arrays.

### LDA Model Training

For model building, test data from centromeres 2, 7, and 10 from the maize NC350 reference genome were used [10]. 229 non-overlapping 10 kb bins were manually labeled as HOR, order, or disorder based on repeat similarity dot plots and network topology. The data was split into training and test sets (n = 183 and 46, respectively). Extracted characteristics were then used as predictor variables.

The first model was an LDA (Linear Discriminant Analysis) model built with MASS (v7.3) [75]. The model uses 3 linear discriminant functions (LD1, LD2, LD3), the first two of which explain 91.33% of the data variation (76.48% and 14.85%, respectively). LD1 utilizes the proportion of monomers collapsed into the most prevalent subtype, proportion of monomers in the second largest cluster, number of unconnected clusters, and proportion of monomers in the largest cluster. LD2 utilizes the proportion of monomers collapsed into the most prevalent subtype, proportion of monomers in the largest cluster, number of unconnected clusters, and the average pairwise Jaccard Index. The LDA model was 89% accurate with the test data.

A second model was also built to compare performance. The second model was a decision tree, built with rpart (v4.1) [76,77]. The decision tree utilized the number of unconnected clusters, modularity, and proportion of monomers in the largest cluster to categorize the bins. This model was slightly less accurate (87%) with the test data.

The LDA model was selected for use due to its better performance in test data. For each bin, erroneous classifications were removed (i.e., if all but one threshold level predicted HOR, the one threshold prediction was removed). Then, the classification prediction for each bin with the highest posterior probability was selected, and tandem bins with consistent classifications were merged.

### HOR Pattern Validation and Purity (Local HOR analysis and HOR purity)

Repeat monomers were identified as described in Identifying repeats in reads and assemblies. For each repeat array, the monomers were extracted from non-overlapping 10-kb bin windows while maintaining their original coordinates within the file structure. Bins with <5 monomers were abandoned. Within a single bin, repeat monomers were compared all-to-all using BLAT and pairwise Jaccard indices calculated (# identical bp/(total length of both sequences - # identical bp)). Jaccard index scores were used to build a set of networks for each bin with thresholds from 0.90 to 0.99 in 0.01 increments.The summary information for each bin was generated as input data for the pre-trained LDA model (see above). The LDA model classified the repeat structures for each bin at every threshold into three classes: HOR, order, and disorder. For each bin, the similarity threshold classified with the highest posterior probability was deemed the “optimal clustering threshold”, and adjacent bins with consistent classifications and thresholds were merged.

To validate identified HOR patterns within a single merged HOR bin, monomer patterns were converted to character strings. First, bins classified as HOR were extracted and monomers were re-clustered based on their optimal clustering thresholds, but without collapsing identical monomers. Monomers were then labeled by their cluster identity. For example, all monomers in the largest cluster were given the label “A”, all the monomers in the second largest cluster were labeled “B”. Characters A-Y, 0-9, and a-z were used as labels. “Z” was used to identify all monomers that exist in a private cluster, meaning the monomer was clustered by itself with no similar sequences.

The character strings were then decomposed into k-mers– starting with k=3 up until k where all k-mers occurred only once. The k-mer list was then filtered based on the initial criteria: all k-mers containing “Z” were removed, k-mers that are majority one letter (like CCCCC or CCAC), smaller k-mers that are fully contained in a larger k-mer that occurs equally often (AB with frequency of 4 removed in favor of ABAB with frequency of 2), and larger k-mers with subsets that occur more often (ABCD with frequency of 3 removed for ABC with frequency of 4 and BCD with frequency of 3). Final k-mer lists contained overlapping patterns to capture both largest HOR sizes and smaller variants and partial patterns. Over 98% of predicted HOR bins were confirmed to have at least 10% content of identifiable HOR patterns of at least 3 monomers, occurring >=2 times within the bin.

For each HOR, purity was then assessed. Purity was calculated as the number of monomers in a string in an identifiable HOR pattern, represented as a k-mer that passed filtering, divided by the total number of monomers in the string. For this value, monomers in overlapping patterns were only counted once. The small number of bins (∼2%) that had no HORs (a purity of 0) were relabeled as order.

### Shared HOR Analysis

To compare HOR patterns among non-adjacent bins or syntenic repeat arrays, consensus representatives of HOR patterns were compared. First, consensus monomers for all monomer subtypes in identified HORs were generated. ClustalOmega (v.1.2.4) was used to make a multiple sequence alignment, and EMBOSS (v6.6) was used to make the consensus [41,78–81]. For example, if the pattern in bin 1 was ABCABCABC, consensus monomers for subtypes A, B, and C were generated.

Then, consensus sequences from bins of the same optimal clustering thresholds were compared using BLAT, as described previously. Bins compared include non-adjacent bins from the same array and bins from syntenic arrays in other inbreds. For example, consensus subtypes A, B, and C from a 3-mer HOR in B73’s centromere 5 may be compared to consensus subtypes J, K, L, and M from a 5-mer in a syntenic array in Mo17, if they had the same optimal clustering threshold of .98. Comparing consensus sequences made it possible to “translate” patterns to identify conserved patterns. For example, in comparing ABC from B73 and JKLM from Mo17, we may find that A and K are at least .98 similar, B and L are at least .98 similar, and C and M are at least .98 similar. From there, we know that ABC and JKLM contain a shared 3-mer, which may indicate recent shared history between the sequences (Figure 5). These values include cases where only a subset of the monomers within an HOR are shared; for instance an HOR might have been composed of ABCD in one array but only ABC is shared in the syntenic array.

Similar to the HOR pattern validation process, similarity matrices of consensus monomers were converted to a network using graph_from_adjaceny_matrix in igraph (v1.2.6) [73]. Edges below the shared optimal clustering threshold were removed and monomers were labeled by cluster identity. Each group was assigned a character as described above. Here, a private monomer, labeled as “Z”, indicates a monomer that does not share homology with another monomer equal to or greater than the similarity threshold. Monomers not included in the shared HOR analysis were also labeled as “Z”. Then, character strings were decomposed into k- mers as described above, but they were not filtered, to allow for identification of shared partial patterns.

Shared HOR analysis was repeated among all knob180 arrays to assess repeating HOR patterns. High-frequency patterns, shared among multiple non-syntenic arrays, and consensus monomers within these patterns were generated.

### Shared HOR Purity

Purity for all HOR bins in Mo17 and CG108 was recalculated using shared HOR patterns at multiple levels— first considering all shared patterns (shared across multiple bins in the same array and/or with at least one other inbred), shared only within maize (present in at least one other maize inbred), and shared with teosinte (present in at least one teosinte). Shared HOR purity was calculated as the number of monomers in a string in an shared HOR pattern, divided by the total number of monomers in the string.

### Whole Array Comparisons

For whole array pairwise comparisons, non-repetitive sequences within arrays were masked using bedtools (v2.29.2) maskfasta [62,63]. Then, syntenic arrays were compared using BLASTn (v2.2.31) (-db {homologs.db} -query {homologs.fasta} -outfmt “6 qseqid sseqid qlen slen length nident pident qstart qend sstart send”) [59]. Pairwise blast hits were merged using bedtools merge, and total base pairs identical were summed. Then, pairwise similarity was calculated in both directions as the total number of identical base pairs, and a Jaccard index calculated from the data ((# identical bp/(total bp of both arrays - # identical bp)).

### Monomer Similarity-to-consensus Visualization

To visualize repeat structure among syntenic arrays, monomers were compared to their consensus. Repeat arrays were extracted from their assemblies using bedtools getfasta. Then, monomers were identified using HMMER nhmmer. The output file (.out) was converted to a bed file. For hits on the opposite strand, start and end coordinates were flipped. Hits were then extracted from the array file using bedtools getfasta. Finally, monomers were compared to the appropriate primary consensus sequence using BLAT (v3.7) [33].

For this comparison, BLAT was utilized because the output format contains convenient match information for conversion into a Jaccard Index. For monomer comparisons, a Jaccard Index is ideal to measure similarity, penalizing for both length and sequence differences. If multiple scores were provided for a hit, only the highest-scoring similarity hit was used. Repeat array structure was represented by plotting each monomer as a dot with the X axis as the genomic coordinate and the Y axis as the Jaccard score to reference in ggplot2 [82].

### Defining Similarity Blocks Within Large Knobs

In the similarity-to-consensus dot plots for some of the largest knobs, a repeating pattern was visible (Figure 6,n7). In these arrays, there are regions of monomers with low similarity to consensus, visible as a vertical stripe in the dot plot, followed by regions of close similarity to consensus, visible as a horizontal line at the the top of the plot. Each unit of this pattern is roughly ∼1 Mb on average. This same striped pattern was observed in the large knob of chromosome 7 in 8 inbreds where the assembled knob was at least 1 Mb (TIL25, K64, CML442, Tx779, Tx777, CG44, CG119 and CG108).

To directly compare the similarity blocks, four 1 Mb windows were selected, three in CG108 K7L and one in CG108 K8L. These windows represent sample similarity blocks. Within each block, monomers were extracted using bedtools (v2.3) getfasta and compared all-to-all with BLAT (v3.7) [30,33], [62,63]. Monomers with at least .98 pairwise Jaccard similarity were represented in a dot plot.

### CENH3 Enrichment

CENH3 illumina CHIP-seq reads and their corresponding input runs from Mo17 were downloaded from NCBI BioProject PRJNA751841 [18]. These reads were generated in the same study as the Mo17 PacBio HiFi reads [18].

For repeat-sensitive mapping, the process from Logsdon et al [4] was used. Briefly, reads were trimmed and dedupped using fastp (--dedup --detect_adapter_for_pe --cut_front --cut_tail) (v.0.23.2) [4,83]. Prepped reads were then aligned to the generated Mo17 assembly using BWA-MEM (v.0.7.17) (-k 50 -c 1000000) [84]. Hits were filtered with samtools (v0.1.19) (view -b -S -F 2308) [68]. To find unique k-mers for sensitive mapping, unique 51-mers were found with meryl (v1.4.1) (meryl count k-51 | meryl equal-to 1 | meryl-lookup -bed-runs) [85]. Unique k-mers were then used to filter alignment bam files, where read alignments that fully overlapped with a unique k-mer were extracted using bedtools (v.2.29) intersect (-b1 {chip.bam} -b2 {unique.bed} -ubam -wa -F 1) [62,63,85]. Uniquely mapping hits were then normalized and 1 kb bins compared using deepTools bamcompare (-of bedgraph --operation ratio --binSize 1000 --scaleFactorsMethod None --normalizeUsing RPKM) [86]. For plotting, CenH3 enrichment was averaged over 100 kb bins.

## Acknowledgements

We thank Arun Seetharam and Matthew Hufford for providing early access to the raw PacBio reads from the TIL lines, and Jonathan Gent, Meghan Brady, Yibing Zeng and Xiao Dong for helpful comments. This study was also supported by resources and technical expertise from the Georgia Advanced Computing Resource Center.

## Peer review information

Andrew Cosgrove and Wenjing She were the primary editors of this article and managed its editorial process and peer review in collaboration with the rest of the editorial team. The peer-review history is available in the online version of this article.

## Authors’ contributions

RDP and RKD conceived and designed the experiments. MCR and ESB provided data for analysis. RDP and MW analyzed the data and visualized it. RDP, MW and RKD interpretated the data. RDP and RKD drafted and revised the manuscript. All authors read and approved the final manuscript.

## Data availability

We used previously published HiFi data for B73 [87], B73-Ab10 [56,88], and HiFi and CENH3 ChiP-seq data from Mo17 [18,89]. Raw PacBio HiFi reads for CG44, CG119, CG108, Tx777, and Tx779 are available at ENA [90]. Raw PacBio HiFi reads for K64 and CML442 are available at ENA [91]. The genome assemblies used in this work are available at Zenodo [92]. All code is available on Github [93] and Zenodo [94]. The downloadable HiReNET tool is available at GitHub [95] and Zenodo [96]. The code is released under the MIT license.

## Declarations

### Ethics approval and consent to participate

Not applicable.

### Consent for publication

Not applicable.

## Additional Files

## Additional file 1: Supplementary Tables

**Table S1:** Proportion of satellite repeat content in older, CLR-based assembly data.

| Line | Data | Total Length (Mb) | CentC (%) | Cent4 (%) | Knob180 (%) | TR1 (%) |
| --- | --- | --- | --- | --- | --- | --- |
| **Ab10** | CLR Assembly | 2,243 | .27% | .011% | 2.8% | .50% |
|  | CLR Pseudomolecules | 2,163 | .19% | .012% | **.059%** | 0.35% |
|  | Illumina 80x | 184,444 | .097% | .0059% | 2.6% | **.020%** |
|  | PacBio CLR 62x | 151,353 | **.035%** | **.0025%** | 1.5% | **.079%** |
|  | ONT 50x | 118,869 | **.032%** | .004% | 1.3% | **.11%** |
| **B73** | Assembly | 2,183 | .22% | .0082% | 1.42% | .15% |
|  | Pseudomolecules | 2,132 | .16% | .0084% | .44% | .15% |
|  | Illumina 21x | 47,778 | .13% | .0071% | 1.8% | .12% |
|  | PacBio CLR 79x | 172,780 | **.054%** | .0029% | .80% | .048% |
| **NC350** | Assembly | 2,291 | .34% | .0079% | 2.7% | 1.17% |
|  | Pseudomolecules | 21,64 | .18% | .0080% | .80% | .16% |
|  | Illumina 69x | 158,744 | .20% | 0.0062% | 4.3% | 1.0% |
|  | PacBio CLR 141x | 323,810 | .088% | .0027% | 1.71% | 0.47% |

Red indicates the value is <75% of the assembly content, bold red indicates the value is <25% of the assembly content. Green indicates the values >125% of assembly content, bold green indicates value is >175% of assembly content.

**Table S2:** Proportion of satellite repeat content in HiFi CCS reads and primary contigs.

| Line | Data | Length (Mb) | CentC | Cent4 | Knob180 | TR1 |
| --- | --- | --- | --- | --- | --- | --- |
| **B73** | CCS (20x) | 48,075 | .046% | .0072% | .37% | .069% |
|  | Contigs | 2,186 | .15% | .0083% | 1.3% | .14% |
|  | Assembly | 2,121 | .067% | .008% | .10% | .13% |
| **Ab10** | CCS (23x) | 53,424 | .038% | .0053% | .74% | .13% |
|  | Contigs | 2,285 | **.15%** | .008% | **3.21%** | **.29%** |
|  | Assembly | 2,133 | .057% | .008% | **.15%** | **.26%** |
| **Mo17** | CCS (66x) | 151,123 | .10% | .006% | .47% | .076% |
|  | Contigs | 2,707 | **.26%** | .005% | **.82%** | .098% |
|  | Assembly | 2,151 | **.32%** | .006% | **1.0%** | .12% |
| **CG108** | CCS (27x) | 62,284 | .33% | .007% | 2.6% | .28% |
|  | Contigs | 2,258 | .34% | .008% | 2.3% | .32% |
|  | Assembly | 2,205 | .35% | .008% | 2.4% | .32% |
| **CG44** | CCS (21x) | 48,968 | .26% | .007% | 2.9% | .10% |
|  | Contigs | 2,249 | .29% | .007% | 2.7% | .11% |
|  | Assembly | 2,217 | .29% | .007% | 3.0% | .11% |
| **CG119** | CCS (17x) | 38,822 | .26% | .006% | 1.9% | .13% |
|  | Contigs | 2,253 | .35% | .007% | **3.4%** | .14% |
|  | Assembly | 2,140 | .37% | .007% | **.36%** | .15% |
| **Tx777** | CCS (21x) | 48,209 | .30% | .007% | 3.6% | .15% |
|  | Contigs | 2,272 | .32% | .007% | 3.3% | .16% |
|  | Assembly | 2,219 | .33% | .007% | 2.9% | .16% |
| **Tx779** | CCS (19x) | 44,744 | .27% | .008% | 4.0% | .91% |
|  | Contigs | 2,315 | .32% | .008% | 4.4% | .97% |
|  | Assembly | 2,276 | .32% | .008% | 4.1% | .81% |
| **K64** | CCS (23x) | 52,074 | .12% | .003% | 1.73% | .44% |
|  | Contigs | 2,333 | .20% | .002% | **5.1%** | .59% |
|  | Assembly | 2,145 | **.22%** | .002% | .47% | .39% |
| **CML442** | CCS (19x) | 43,183 | .28% | .008% | 3.43% | .71% |
|  | Contigs | 2,413 | .33% | .008% | 5.20% | .67% |
|  | Assembly | 2,255 | .35% | .007% | 3.0% | .69% |
| **TIL01** | CCS (28x) | 76,038 | .27% | .012% | 6.4% | 1.8% |
|  | Contigs | 2,766 | .47% | .010% | 8.9% | 1.9% |
|  | Assembly | 2,540 | **.50%** | .010% | 9.4% | 1.9% |
| **TIL11** | CCS (21x) | 54,395 | .20% | .007% | 2.1% | 1.32% |
|  | Contigs | 2,459 | **.48%** | .007% | **4.8%** | 1.55% |
|  | Assembly | 2,263 | **.44%** | .007% | 1.5% | 1.5% |
| **TIL25** | CCS (25x) | 55,175 | .33% | .003% | 2.3% | .14% |
|  | Contigs | 2,134 | **.67%** | .002% | 3.3% | .14% |
|  | Assembly | 2,085 | **.66%** | .002% | 3.4% | .14% |

Red indicates the value is <75% of the CCS content, bold red indicates the value is <25% of the CCS content. Green indicates the values >125% of CCS content, bold green indicates value is >175% of CCS content.

**Table S3:** Contig Counts and Assembly N Gaps.

| **Line** | **Contigs** | **Contigs in**<br>**Assembled**<br>**Chr** | **N Gaps in**<br>**Assembly** | **N Gaps**<br>**in Arrays** |
| --- | --- | --- | --- | --- |
| **B73** | 1248 | 136 | 153 | 18 |
| **Ab10** | 2117 | 162 | 148 | 16 |
| **Mo17** | 8795 | 56 | 46 | 0 |
| **CG108** | 774 | 57 | 72 | 2 |
| **CG44** | 730 | 88 | 101 | 1 |
| **CG119** | 1074 | 154 | 162 | 5 |
| **Tx777** | 910 | 217 | 224 | 11 |
| **Tx779** | 777 | 277 | 288 | 11 |
| **K64** | 1259 | 78 | 93 | 11 |
| **CML442** | 1580 | 124 | 141 | 9 |
| **TIL01** | 6304 | 176 | 166 | 12 |
| **TIL11** | 2113 | 131 | 121 | 16 |
| **TIL25** | 906 | 74 | 64 | 6 |

**Table S4:**
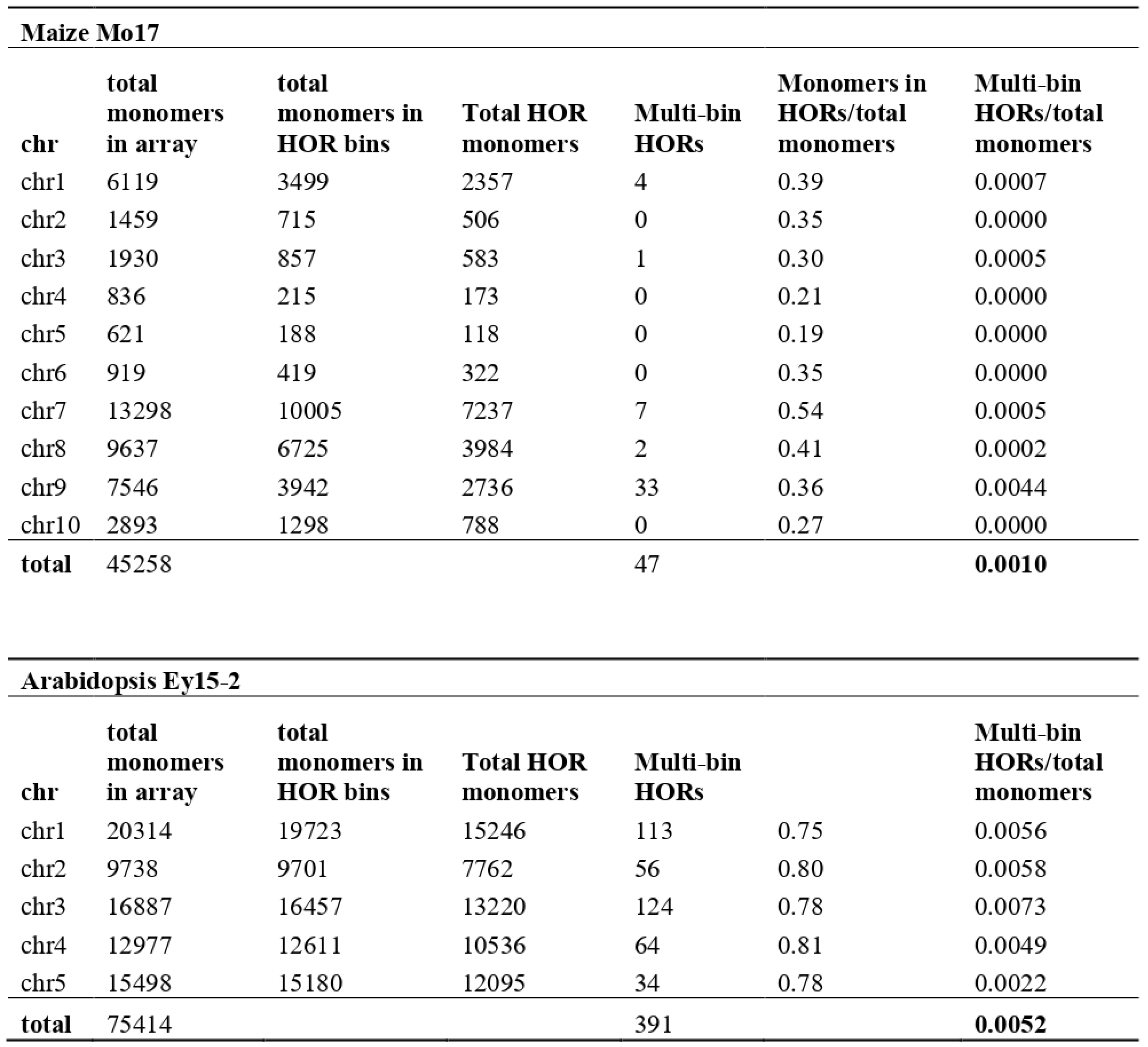
HORs and multi-bin HORs in Maize Mo17 and Arabidopsis Ey15-2 centromeres.

**Table S5:** Average syntenic array similarity as measure by Jaccard index. The “no N gaps column” shows values from Figure 3e (y axis) as total averages. Also shown below are the equivalent values when considering all arrays including those with N gaps.

| **Satellite** | **# Arrays** | **Mean**<br>**Jaccard**<br>**Similarity** | **Mean**<br>**Jaccard**<br>**Similarity,**<br>**no N gaps** |
| --- | --- | --- | --- |
| **All** | 95 | 0.82 | 0.82 |
| **Cent4** | 1 | 0.96 | 0.96 |
| **CentC** | 18 | 0.86 | 0.88 |
| **knob180** | 64 | 0.82 | 0.82 |
| **TR1** | 12 | 0.74 | 0.74 |

**Table S6:** HOR patterns from Mo17 and CG108 that are observed in at least one other syntenic array.

| **Inbred** | **Satellite** | **# HORs in analysis** | **# HORs that are shared** | **% of HORs that are shared patterns** |
| --- | --- | --- | --- | --- |
| **Mo17** | **All** | 1470 | 200 | 14% |
|  | **Cent4** | 8 | 7 | 88% |
|  | **CentC** | 342 | 46 | 13% |
|  | **knob180** | 1006 | 76 | 8% |
|  | **TR1** | 114 | 71 | 62% |
| **CG108** | **All** | 3343 | 637 | 19% |
|  | **Cent4** | 8 | 7 | 88% |
|  | **CentC** | 416 | 53 | 13% |
|  | **knob180** | 2512 | 335 | 13% |
|  | **TR1** | 407 | 242 | 59% |

**Table S7:** Knob High-Frequency HORs.

| **Pattern** | **Total**<br>**Frequency** | **N**<br>**regions** | **N freq in**<br>**K6L** | **N freq in**<br>**K7L** | **N freq in**<br>**K8L** | **Mean**<br>**Distance**<br>**Between**<br>**Regions** | **Mean**<br>**Frequency**<br>**within**<br>**each**<br>**Region** |
| --- | --- | --- | --- | --- | --- | --- | --- |
| **ADBA** | 102 | 33 | 0 | 65 | 37 | 1,163,183 | 4.5 |
| **AADB** | 102 | 32 | 0 | 52 | 50 | 1,473,366 | 5.1 |
| **BAAD** | 85 | 30 | 2 | 48 | 35 | 1,473,366 | 4.1 |
| **ADB** | 348 | 67 | 0 | 181 | 167 | 677,293 | 7.7 |
| **ADC** | 120 | 36 | 8 | 63 | 49 | 1,146,341 | 6.5 |
| **DBA** | 141 | 46 | 4 | 85 | 52 | 953,519 | 5.8 |
| **DBG** | 116 | 30 | 2 | 61 | 53 | 1,227,805 | 4 |
| **DBEF** | 70 | 30 | 0 | 41 | 29 | 1,481,578 | 4.8 |
| **BEF** | 87 | 36 | 2 | 53 | 32 | 1,185,263 | 4.6 |
| **DBE** | 113 | 40 | 0 | 63 | 50 | 1,128,822 | 5.8 |

**Table S8:**
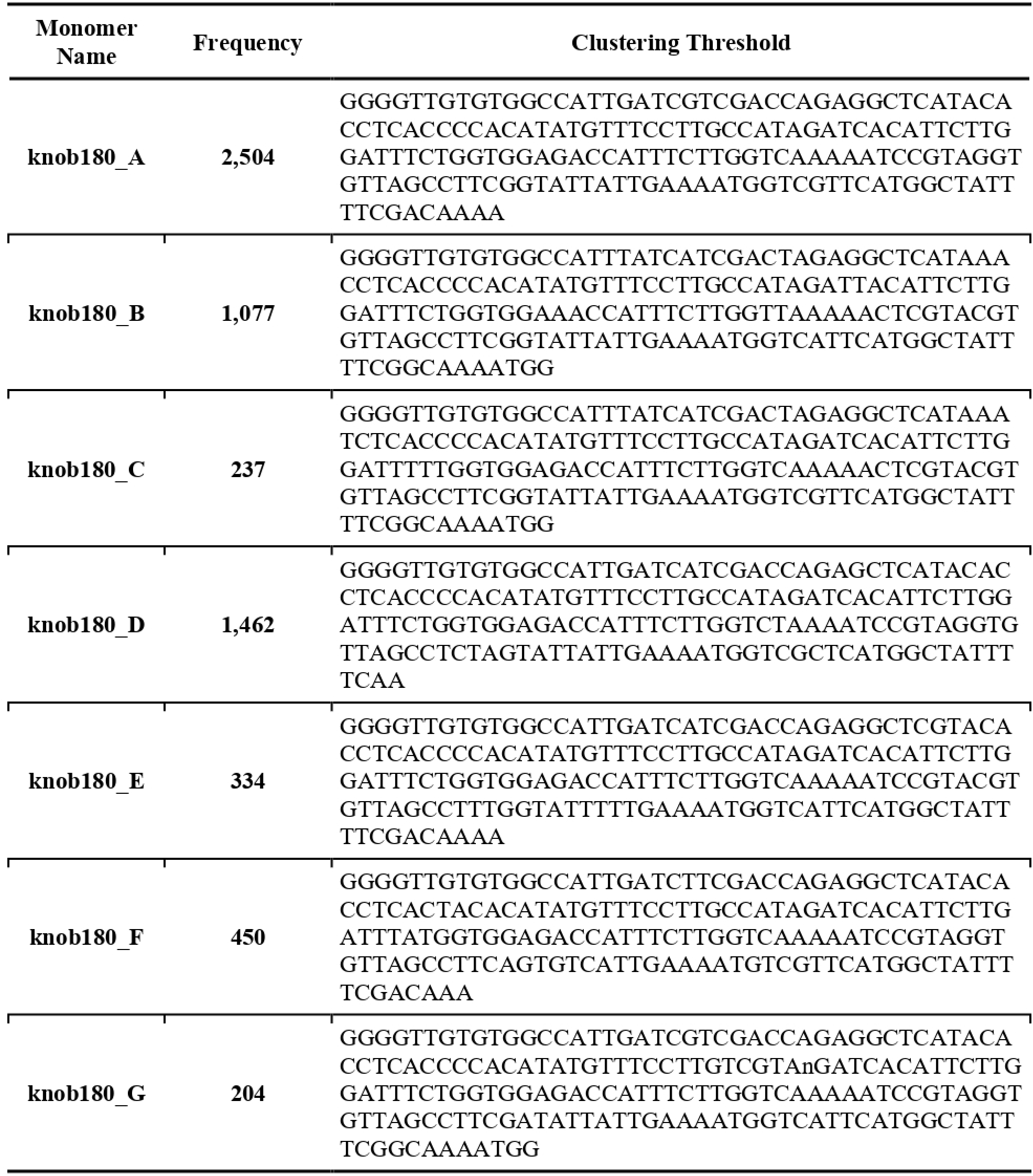
Knob High-Frequency HOR Monomers.

## Additional file 2: Supplementary Figures

**Fig. S1.**
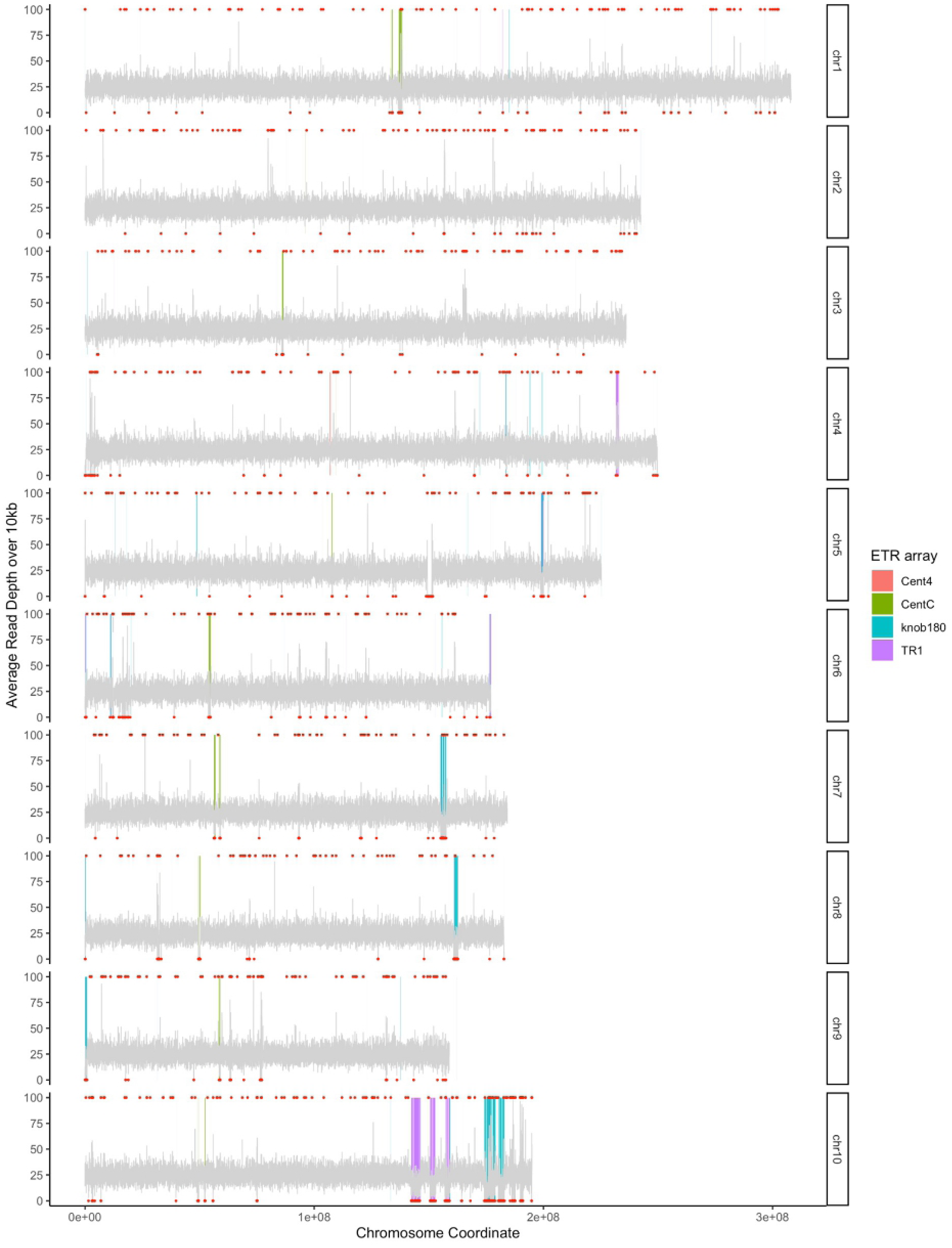
PacBio HiFi reads aligned to the previously published B73-Ab10 reference assembly. Grey lines represent 10 kb average read depth. Red dots above and below make 10 kb bins where the average read depth falls outside two standard deviations of the mean depth. Satellite arrays are represented by colored boxes. Note that satellite repeat areas appear to have lower coverage than that other parts of the genome. This is likely because the repeat areas were overpolished, such that the more accurate HiFi reads do not align well (see also Fig. S2).

**Fig. S2.**
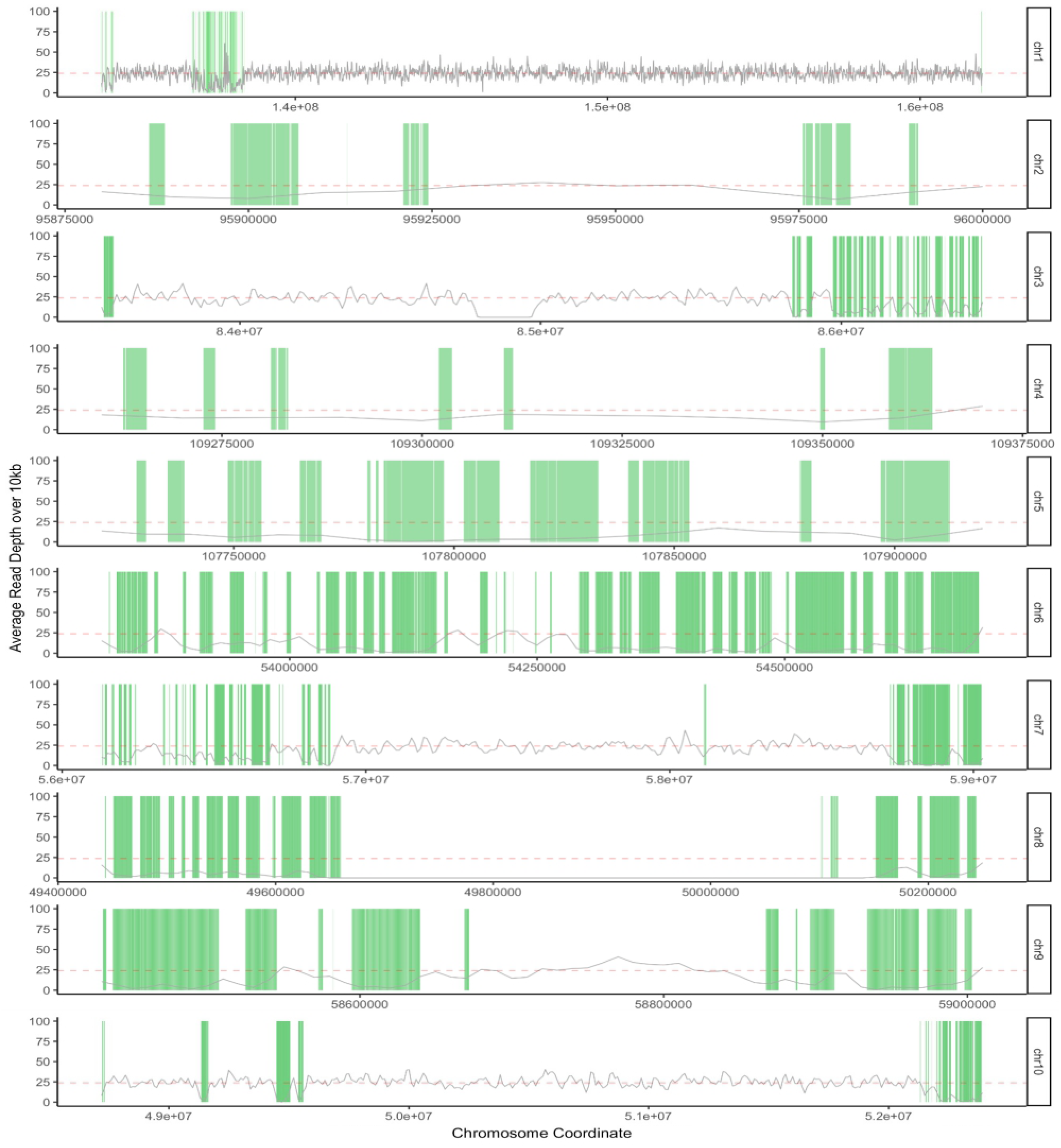
PacBio HiFi reads aligned to the CentC regions of the previously published B73-Ab10 reference assembly. This reference was assembled using older CLR technology. There is one region of high coverage in the CentC area of chromosome 1 (suggesting over-collapsing during assembly), but, overall, unexpectedly poor alignment of HiFi reads over CentC-rich areas. Grey lines represent 10kb average read depth. Red dotted line represents the average read depth. CentC satellites are represented by green boxes.

**Fig. S3.**
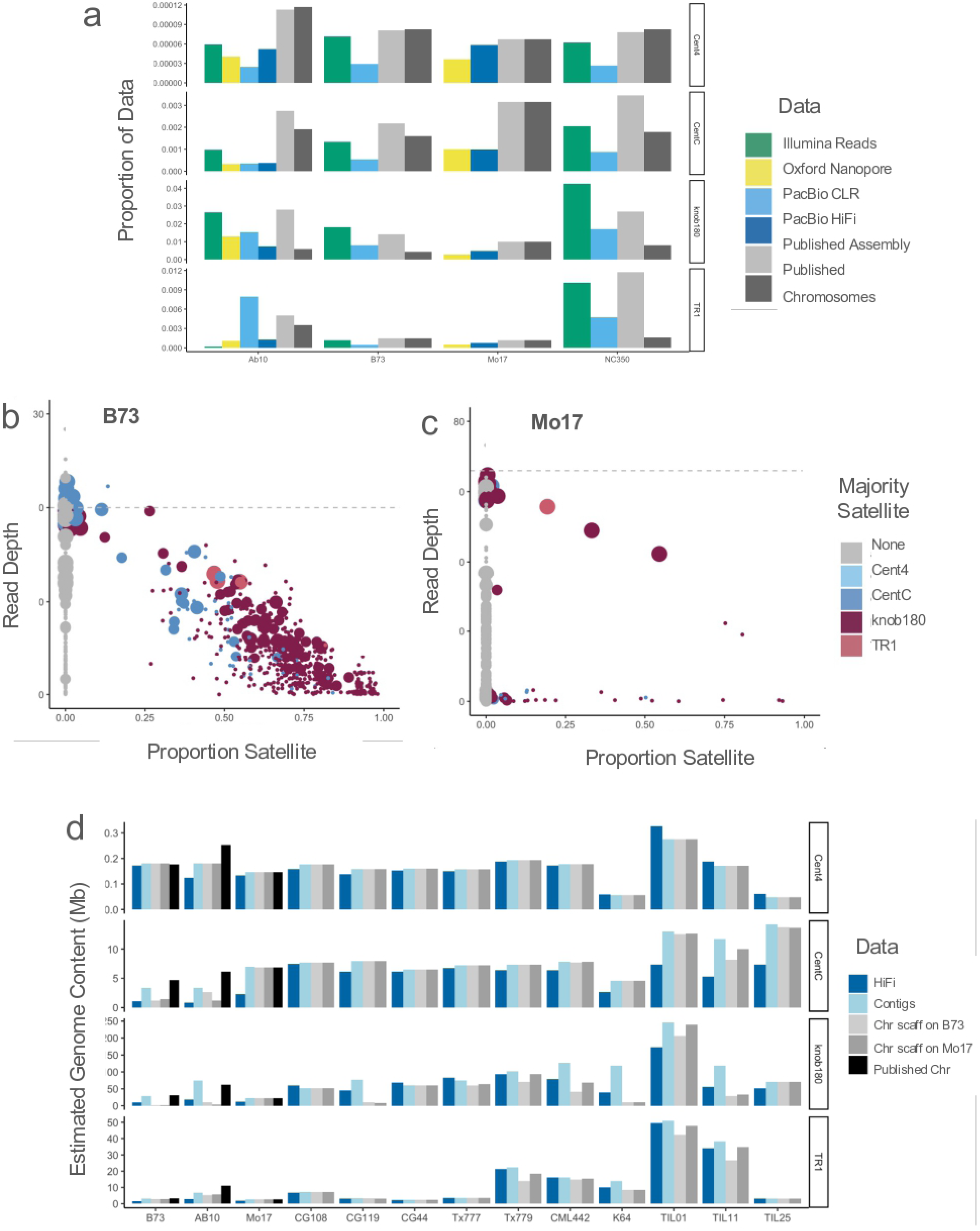
Assembly Repeat Biases. **a**) Satellite content of raw reads and assembled genomes from four previously-assembled inbred lines. **b**) Hifiasm-generated contig read support and satellite content for B73. Size represents contig size. **c**) Hifiasm-generated contig read support and satellite content for Mo17. **d**) Satellite content of PacBio HiFi reads and hifiasm-generated contigs for 13 inbreds, compared to previously-published genome assembly and pseudo-molecules generated from the high-confidence set of hifiasm-generated contigs scaffolded to B73 and Mo17 (i.e., after contigs with lower than half the expected read support were removed).

**Fig. S4.**
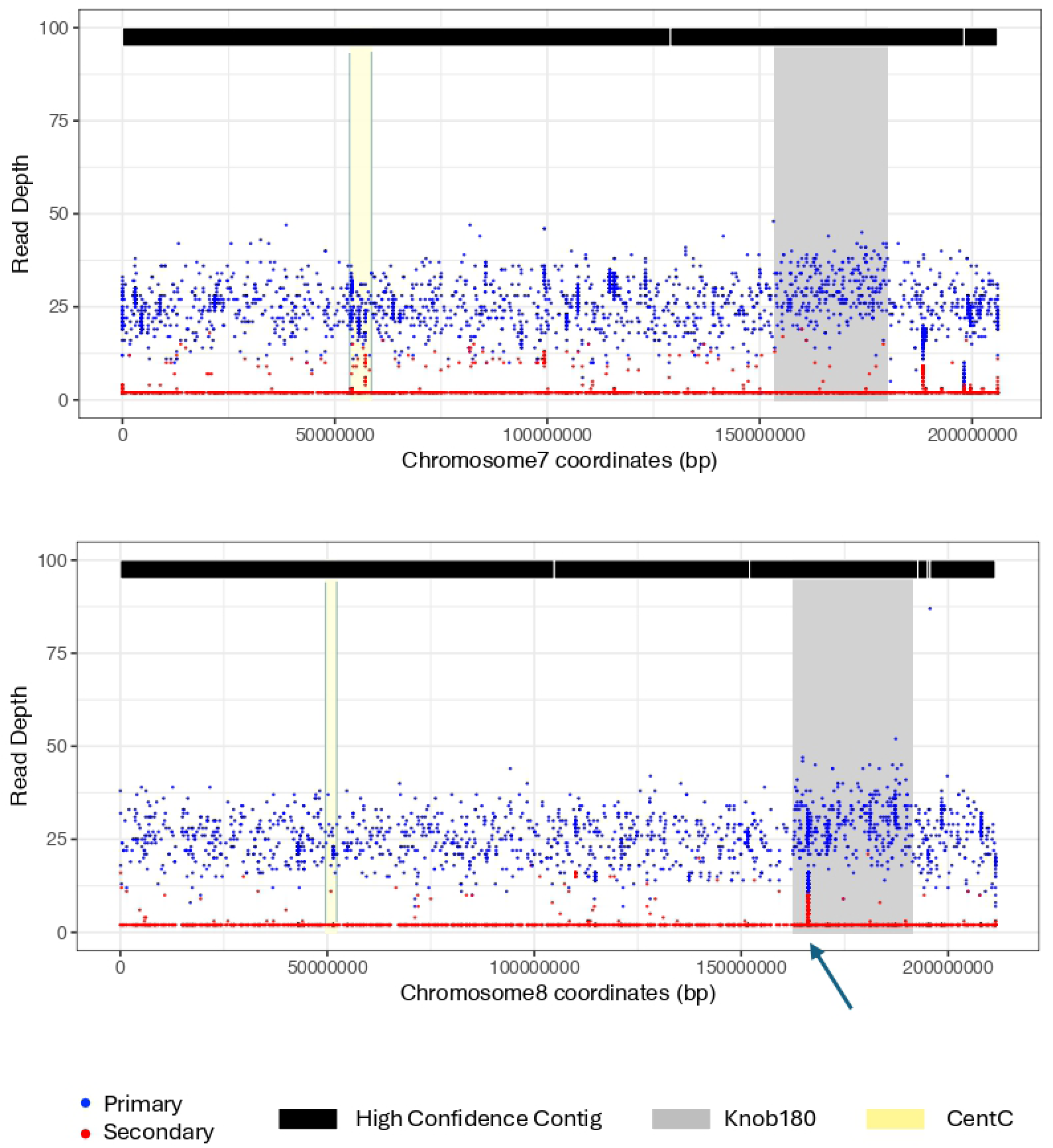
Primary and secondary alignments of raw HiFi reads from CG108 to the CG108 assembly of chromosomes 7 and 8. The black blocks at the top show high confidence contigs within the chromosomes (gaps in the black blocks are assembly gaps). The knob180 and CentC regions are indicated. The blue and red dots represent the primary and secondary read counts. The relatively even overall coverage and low frequency of secondary alignments across repeat regions indicates that most reads from repeat arrays are properly assembled. There may be a small area of mis-assembly on the large knob of chromosome 8 (arrow).

**Fig. S5.**
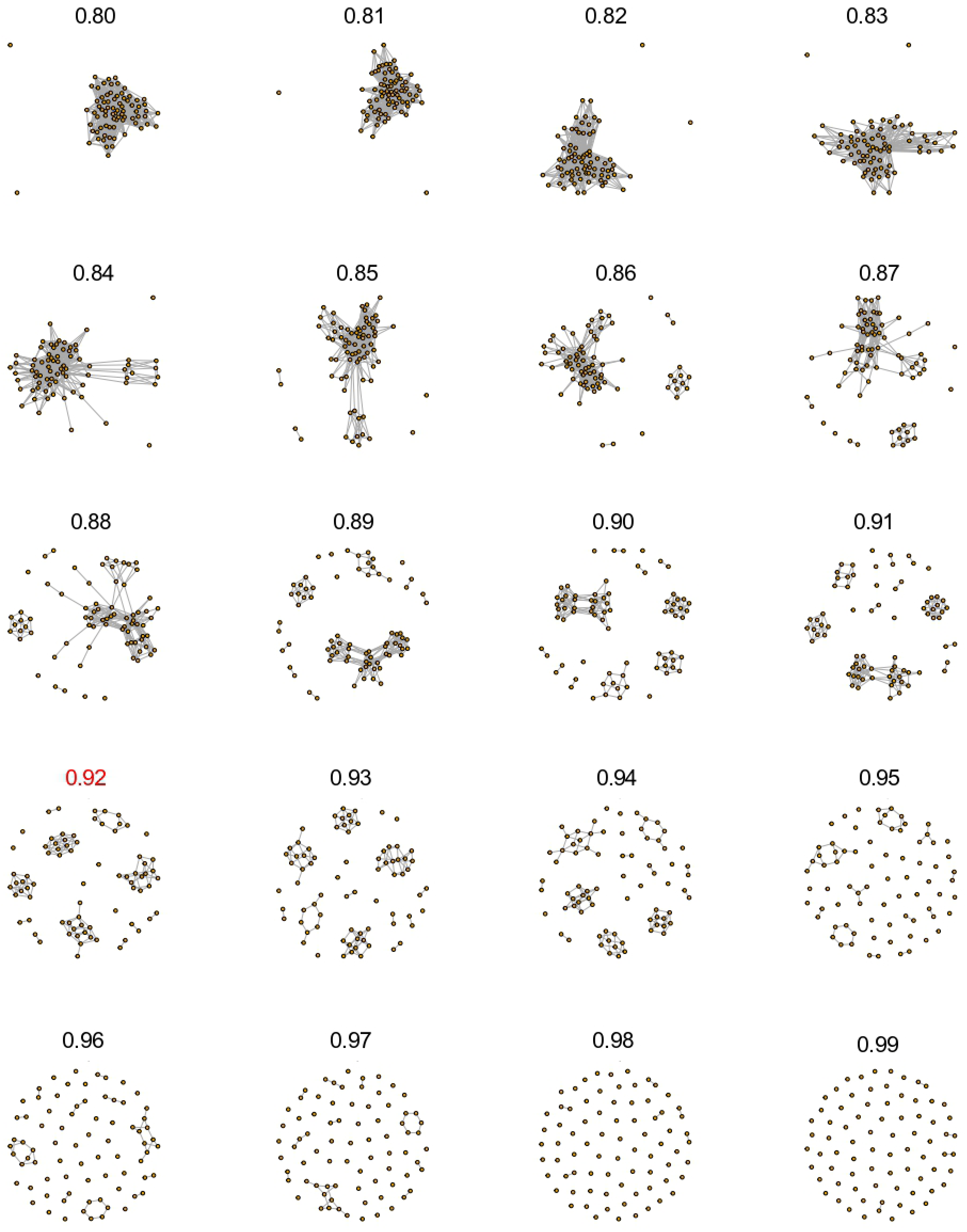
CentC network plots showing clustering thresholds ranging from 0.80 to 0.99. Each node represents a CentC monomer. Note that monomers tend to collapse into one cluster at low thresholds. The range used in HiReNET is 0.90-0.99. The optimum clustering threshold for this bin is 0.92. These data are from the NC350 inbred.

**Fig. S6.**
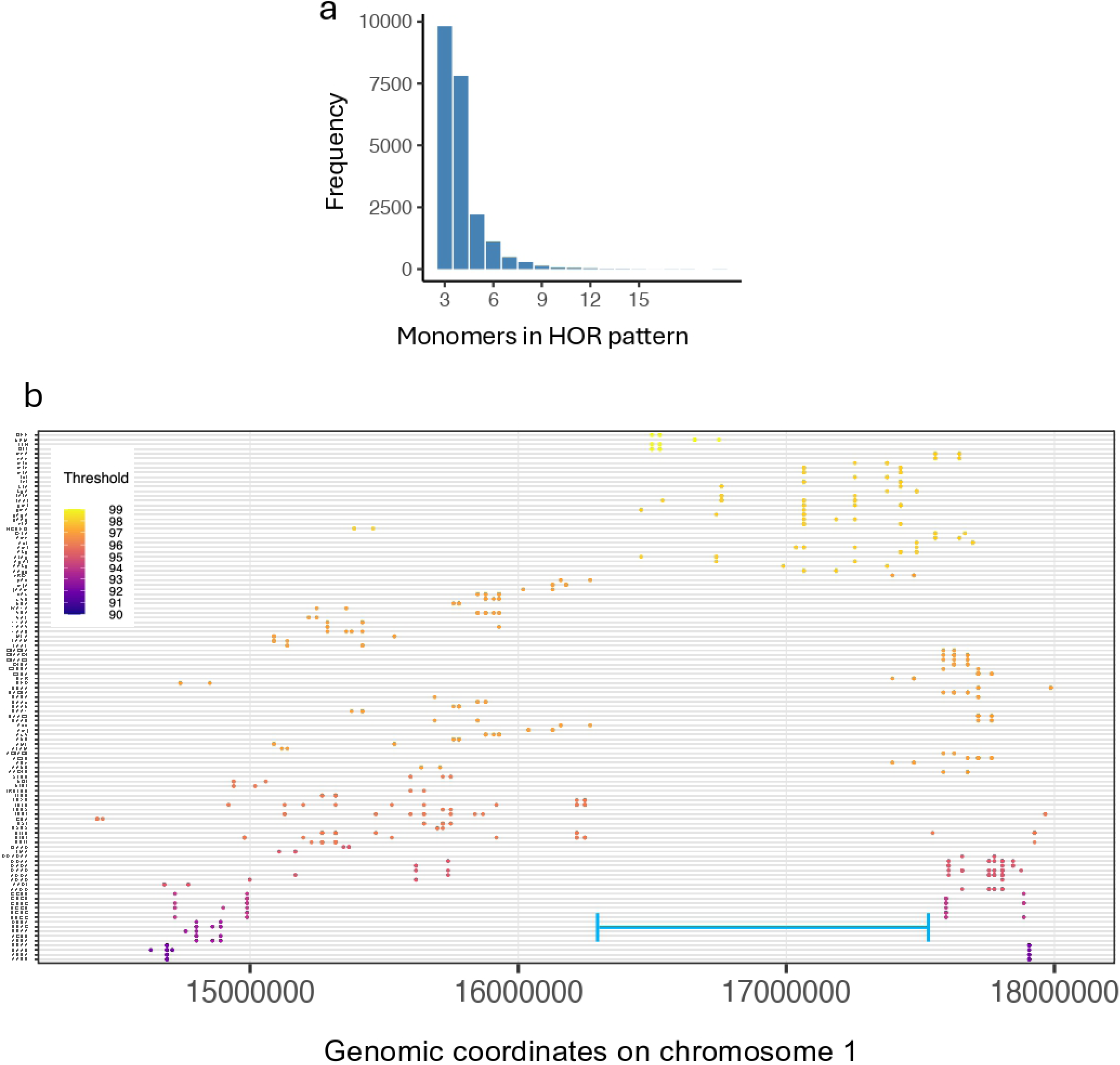
AthCEN178 HOR characteristics and multi-bin AthCEN178 HORs as assayed using HiReNet. **a**) Frequencies of AthCEN178 HOR sizes in the Arabidopsis Ey15-2 ecotype. **b**) Multi-bin AthCEN178 HOR locations within centromere 1 of Ey15-2. The y-axis shows HOR patterns that are found in two or more bins, and the x-axis shows coordinates on centromere 1. The blue bar within the image indicates the region of high HOR score noted in Wlodzimierz et al. 2023.

**Fig. S7.**
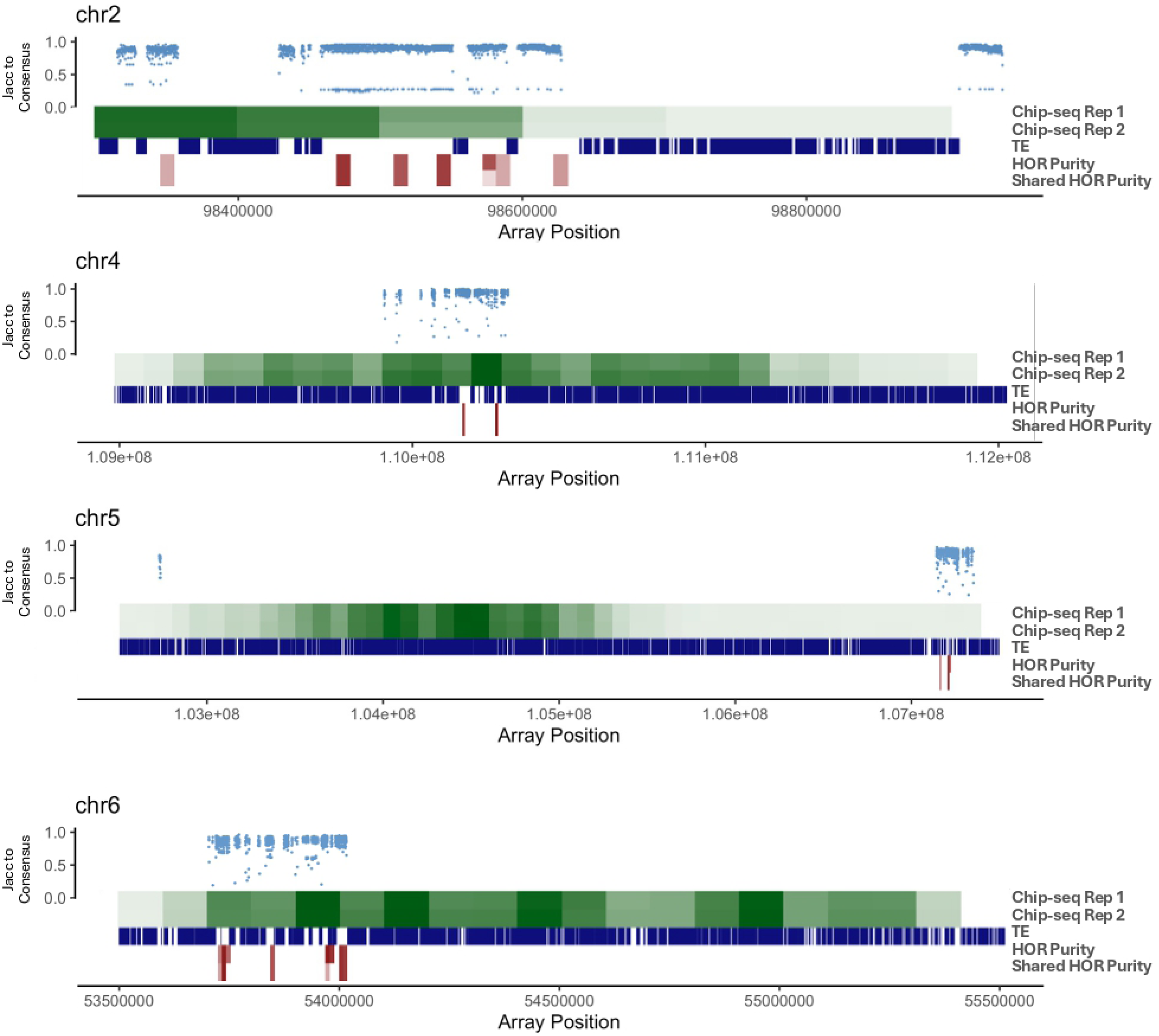
CENH3 Chip-seq on Mo17. Centromere structure of chromosomes 2, 4, 5, and 6. Relative ChIP-seq density in 100 kb bins represented by shades of green. TE presence is represented by blue boxes. HOR purity and shared HOR purity within bins are represented in red. Darker shades of red represent higher purity values.

**Fig. S8.**
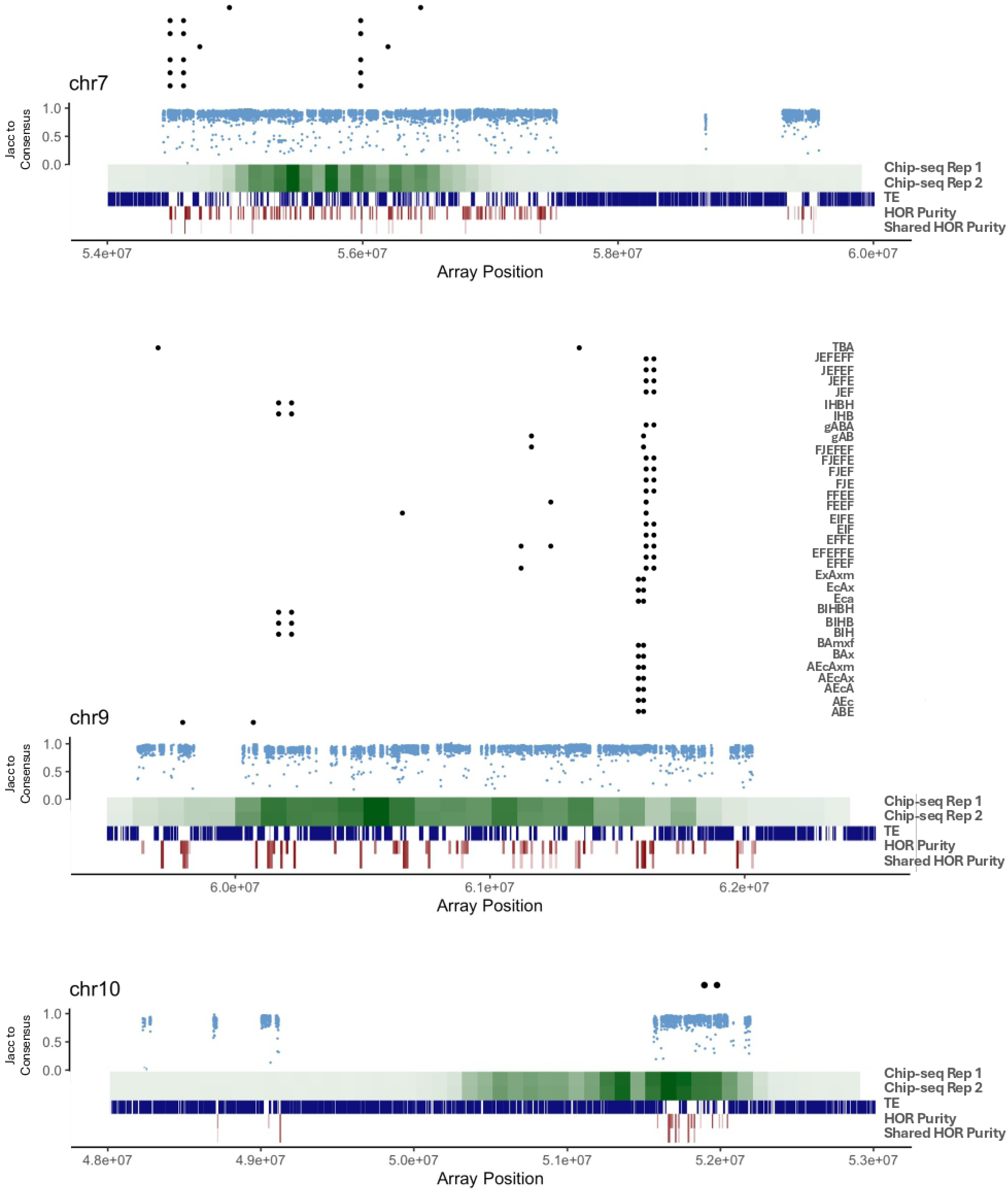
CENH3 Chip-seq on Mo17. Centromere structure of chromosomes 7, 9, and 10. Shared HOR patterns that occur in at least two distinct HOR regions. Dot indicates presence in array, x axis is genomic position. Relative Chip-seq density in 100 kb bins is represented by shades of green. TE presence is represented by blue boxes. HOR purity and shared HOR purity within bins are represented in red. Darker shades of red represent higher purity values.

